# A century of allopatry: plasticity and rapid selection shape phenotypic trait variability under contrasting environments

**DOI:** 10.1101/2025.06.26.661825

**Authors:** Hervé Rogissart, Martin Daufresne, Guillaume Evanno, Jean Guillard, François-Raphael Lubin, Emilie Chancerel, Allan Raffard

## Abstract

Allopatric isolation under contrasting environments can drive rapid phenotypic divergence, even over contemporary timescales. Rapid changes in morphology or physiology can allow organisms to adapt to biotic and abiotic characteristics of their habitats. While studying metabolism, growth and resources needs may allow to understand adaptation to several selective pressures, these traits are rarely jointly considered. We investigated morphological, growth, and metabolic divergence in two allopatric populations of Arctic charr (*Salvelinus alpinus*) sharing a common evolutionary origin but inhabiting contrasting environments. We combined field observations, common garden and quantitative genetic approaches to disentangle contributions of genetic divergence and plasticity to phenotypic variability. Wild adults differed in body shape and growth trajectories, potentially reflecting plasticity related to resource availability and temperature variations. Under common garden conditions, juveniles displayed inter-population differences in routine metabolic rate, its allometric scaling with body mass. These patterns suggest divergent selection on physiological traits. Despite low neutral genetic differentiation, phenotypic divergence unfolded in fewer than 100 years, suggesting that plasticity and selection can promote rapid multi-trait changes. These findings highlight that considering changes in physiological, growth and morphological traits can reveal the adaptive potential of small, isolated populations facing rapid environmental change.

## 1. Introduction

Phenotypic variation and organismal performance shape how populations cope with environmental challenges, persist over time and evolve under selective pressures. Populations can respond to environmental variation through rapid phenotypic changes, classically studied through the lens of phenotypic plasticity [1]. Phenotypic plasticity is defined as the ability of a single genotype to produce different phenotypes in response to varying environmental conditions [1]. When adaptive, it enables populations to maintain their fitness, through a range of physiological, morphological and behavioural processes [2]. This capacity is notably illustrated by rapid changes in morphological traits, such as head size and shape in green anole (*Anolis carolinensis*), which enhance feeding performance under novel trophic conditions [3]. It also includes transgenerational plasticity in physiological traits, such as metabolic rate, which supports growth and energy balance under changing temperatures and food sources [4]. The ability of individuals to adjust their phenotypes to novel environmental conditions can vary among individuals and populations and is determined by complex interactions between genetic and environment, thereby shaping inter-population phenotypic variability [5]. While phenotypic plasticity can operate within a generation and across generations through transgenerational plasticity, rapid evolution can also shape rapid phenotypic differentiation among populations within just a few generations [6].

The rapid evolution of phenotype traits often occurs in populations exposed to contrasting environments, whether across space or time, where local selective pressures drive adaptive responses [6,7]. However, it may also arise through non-adaptive processes such as genetic drift [8] or from anthropogenic selective pressures [9]. Such rapid evolutionary divergences have been documented in a wide range of taxa, including plants [10], amphibians [11], birds [12] and fish [13]. Studies on rapid evolution have shown that adaptive phenotypic changes can occur within just a few generations, particularly in species facing strong selection pressures [13,14]. For instance, in salmonids, traits such as growth, metabolism and thermal tolerance can evolve in response to environmental variation in 10 to 20 generations [7,15]. Studying these rapid changes on single traits allow to answer precise ecological hypotheses, such as -amongst others-the determinants of the benthic-limnetic morphology of fish or the thermal adaptation of metabolic rate [4,16]. However, since evolutionary divergence often involves complex interactions among numerous traits, a full understanding of adaptation to temperature regimes requires considering coordinated changes of traits linked to energy acquisition, allocation, and use [17].

Many organisms, such as ectotherms, exhibit diverse phenotypic responses to environmental variability, provided by changes in physiology, morphology, behaviour, and life-history traits [18,19]. Metabolic rate is one of the most important traits for the survival and performance of individuals because it is directly linked to energy allocation, determining individual ability to grow, reproduce and tolerate changing temperature fluctuations [20,21]. Metabolism is indeed dependent of genetic and environmental effects [22,23], and determines the pace of energy turnover and physiological individual performance (sometimes referred to as functional traits ; [19,20]). Accordingly, when the temperature rises, the metabolic rate generally increases due to increased enzymatic activity and higher oxygen demand, whereas it decreases when temperature drops, leading to reduced energy expenditure [19,21]. Complementary dimensions of phenotypes, such as morphological traits in fish, are important in mediating resource acquisition and habitat exploitation. Together, changes in metabolism and morphology can allow organisms to respond to variation in temperature or prey availability [4,17,23]. Since morphology and metabolism underlie multiple functions, their variation due to environmental conditions often affects growth rate. Growth indeed reflects the energetic allocation to tissue production [19], which is linked to resource-use efficiency and metabolic demands [16]. Therefore, growth rate likely reflects the cumulative effects of metabolism and resource use, providing a proxy of individual fitness and population performance [25]. Given the thermal dependence of physiological traits, considering joint changes in energetic needs, through metabolism, morphology and allocation to growth is important to fully understand adaptation to thermal environments [2,19,23].

Many valuable insights regarding the mechanisms of rapid evolutionary divergence have emerged from studies focusing on introduced populations, with numerous examples in fish species, such as salmonids [7,26]. For instance, Hendry *et al*. [7] demonstrated the emergence of reproductive isolation in fewer than 13 generations between two sockeye salmon (*Oncorhynchus nerka*) populations that colonized distinct spawning environments, resulting in divergence in adult body morphology and embryonic development traits. Similarly, Koskinen *et al*. [26] documented strong directional selection on early life-history traits (e.g., yolk-sac volume, growth rate) among grayling (*Thymallus thymallus*) populations originating from a single introduction event, despite severe bottlenecks and very low effective population sizes. Moreover, several cases of rapid divergence, both in sympatry [27] and in allopatry [28], have been documented in Arctic charr (*Salvelinus alpinus*). This species is known for its high phenotypic variability and polymorphism [29]. Furthermore, as a cold stenothermic species, the Arctic charr is particularly sensitive to thermal variations, making it a valuable model for studying how environmental pressures influence phenotypic differentiation, plasticity and evolution [30]. Many populations have been introduced at the southernmost edge of the species’ natural distribution, such as in the French Alps [31]. These edge-of-range populations provide unique opportunities to study rapid phenotypic differentiation [32]. These environments often expose populations to suboptimal or novel selective conditions including warmer temperatures, shorter growing seasons and reduced habitat productivity, that differ from core habitats [33]. As such, marginal populations are important for understanding species’ responses to environmental change, particularly under ongoing climate warming [34]. These edge-of-range populations are shaped by both environmental variation and demographic constraints, and their interaction modulates the evolution and expression of plasticity [32,33]. Studying the mechanisms of adaptation and divergence among these populations is essential to assess species capacity to respond rapidly to rapid environmental change [2].

In this study, we aimed at testing the drivers of phenotypic differentiation between two allopatric Arctic charr populations sharing a recent common origin. We focused on populations in Lake Geneva (where Arctic charr is native) and in Lake Allos, into which Arctic charr were introduced from Lake Geneva about a century ago (Figure 1). Using comparative analyses (*Q*_*st*_ − *F*_*st*_ or *P*_*st*_ − *F*_*st*_) we aimed at disentangling the relative contributions of adaptive (e.g., selection) and neutral processes (e.g., genetic drift) to phenotypic differentiation. We first tested whether adults (F_0_) differed in their morphology and growth rate. Lake Geneva is a large, productive and warmer lake than Lake Allos (Table S1), it should therefore provide higher resource availability and longer growing seasons resulting in individuals with higher growth rate compared to individuals from Lake Allos [35–37]. Since morphology is supposedly linked to resource acquisition in fishes [16], differences in competition with other species (i.e., Lake Geneva hosts a diverse fish assemblage, while Lake Allos is hosting a fish community mainly composed of salmonids, Table S1), and in the type of resources between lakes is also expected to generate morphological differentiation higher than expected by genetic drift only (*P*_*st*_ > *F*_*st*_). Second, we tested the genetically based difference of phenotypic variability (metabolism, morphology, growth) using F_1_ individuals raised in a common garden. The metabolic rate was measured at different temperatures (6°C and 10°C) to assess differences in thermal plasticity between populations. Individuals from harsher environments (here Lake Allos) are theoretically expected to show higher performance than individuals from other populations [38]. Accordingly, since growing season and resource abundance are lower in Lake Allos (Figure S1 & Table S1), we expected juveniles from Allos to exhibit higher metabolic rate, plasticity and growth rate, suggesting that selection might alter physiological rate (*Q*_*st*_ > *F*_*st*_). Complementarily, controlled feeding conditions in common garden should result in limited morphological differentiation among populations (*Q*_*st*_ ≈ *F*_*st*_), given that morphology is partly plastic in fish [39]. Comparison of trait differentiation between F_0_ and F_1_ individuals is interesting at it provides insights into the relative role of plasticity and selection in shaping phenotypic differentiation.

**Figure 1.**
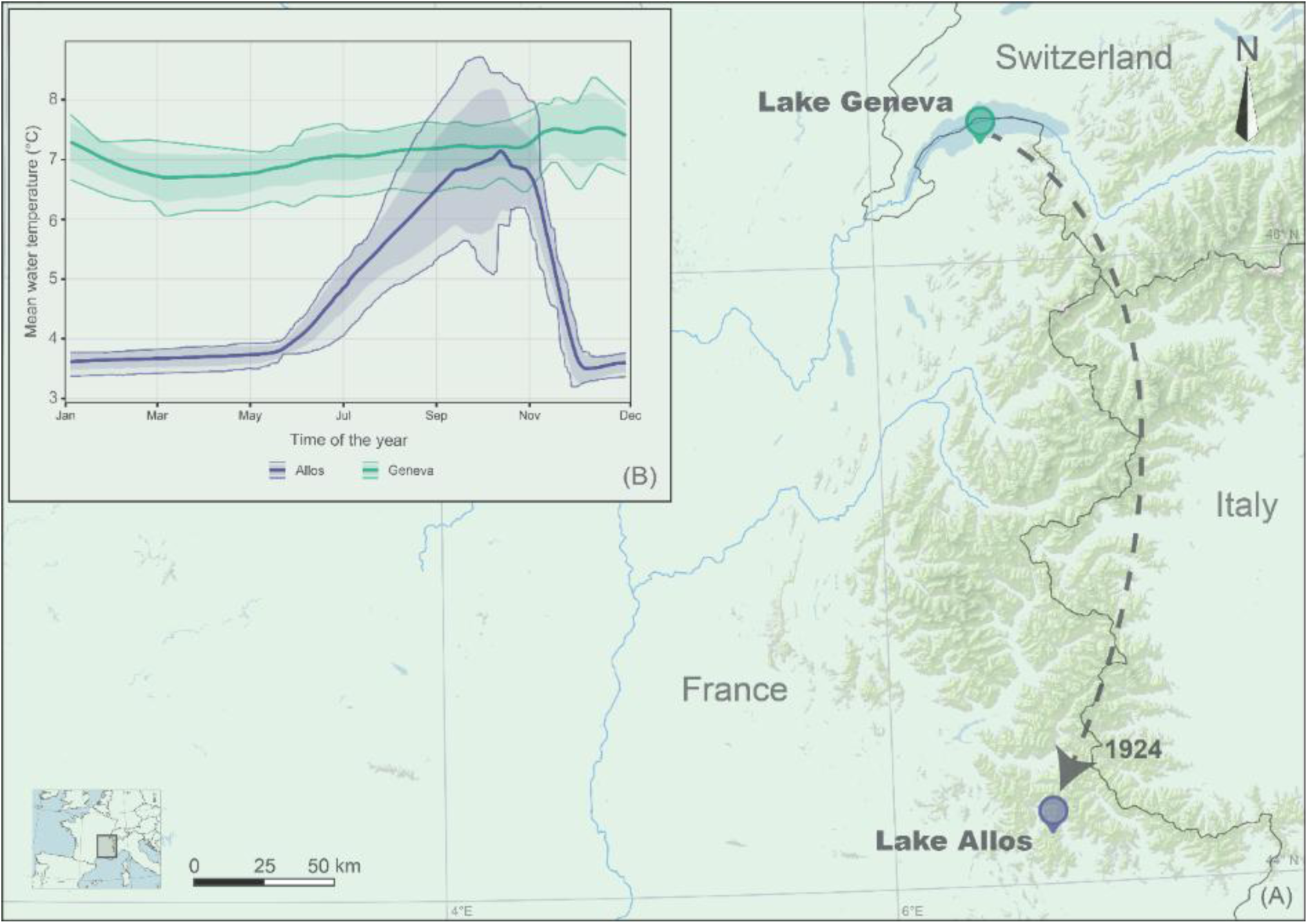
(A) Location of sampling site situated at the southern edge of the species’ distribution range. The most recent documented introduction of Arctic charr (*Salvelinus alpinus*) from Lake Geneva into Lake Allos occurred in 1924 [42]. (B) Mean annual water temperature (°C) for Lake Allos (10–30 m depth, blue) and Lake Geneva (30–100 m depth, green), averaged over 2016–2022, corresponding to the typical depth range inhabited by Arctic charr in each lake. Solid lines represent mean daily temperatures, fine solid lines indicate minimum and maximum values, and dark shaded areas indicate the 95% confidence interval. The map background indicates elevation (m) above 1500 m. Digital elevation model data from [83]. Data were collected for lake Allos from Réseau National de suivi de la Température des plans d’eau, coordinated by pôle R&D ECLA (OFB–INRAE–USMB) and for Lake Geneva from OLA-IS [84], AnaEE-France, INRAE, Thonon-les-Bains and CIPEL.

## 2. Materials and methods

### 2.1 Model species

Arctic charr (*S. alpinus*, Linnaeus, 1758) is the northernmost freshwater fish species, primarily associated with boreal and circumpolar regions [29]. Its native range reaches its southernmost limit in the populations of deep lakes within the French peri-Alpine arc [31]. This cold-adapted salmonid typically spawns in late autumn on coarse substrates, with eggs developing over winter and juveniles emerging in spring [35]. Individuals usually reach sexual maturity from 3 years old [40]. Due to its high phenotypic variability, Arctic charr is considered a species complex [41]. The populations of Arctic charr in the Alps are sometimes referred to as Alpine charr (*S. umbla*) [41].

### 2.2 Study sites

This study focuses on two contrasting populations of Arctic charr from Lake Geneva and Lake Allos (Figure 1, Table S1). Reproduction occurs between November and January, when water temperature ranges from 4°C to 7°C, which are considered optimal for spawning [30,35]. During this period, mean water temperature at spawning depth (10-20 m and 40-60 m for Allos and Geneva, respectively) was 4.2 ± 1.1°C in Lake Allos and 7.9 ± 0.7 °C in Lake Geneva (Figure S1; Table S1). Mortality during egg development increases sharply when incubation water temperatures exceed 9°C, with thermal tolerance reaching its upper limit above 12°C [35]. Arctic charr were introduced from Lake Geneva to Lake Allos through documented stockings in 1922 and 1924 [42]. Overall, this population has not been subjected to any other stocking practices since 1924 and has therefore evolved independently for a century [42]. The contrasting environmental conditions (e.g., thermal regimes, trophic status, resources availability) between these two lakes make them a valuable natural setting for investigating rapid phenotypic differentiation and assessing evolutionary processes shaping trait divergence.

### 2.3 Sampling and rearing in a common garden experiment

A total of 28 adults from Lake Allos were captured through scientific sampling, and 22 from Lake Geneva by professional fishers. In both cases sexually mature individuals were captured on spawning grounds using gillnets (mesh 27–50 mm) during the spawning season in late November 2023 (Table S2). This sampling targeted the sexually mature individuals from the spawning stock of the lake. Eggs from each female were collected and fertilized with milt from a single male of the same origin population (Allos or Geneva) on November 24, 2023 (Table S2). In total, eight full-sib families were established for the Allos population and six for the Lake Geneva population. Hatching dates were similar between populations, and accumulated degree-days (ADD; i.e. cumulative sum of daily mean water temperatures since fertilization) to hatching were comparable (Table S3). Only these independent full-sib families were used in all quantitative analyses; whereas additional half-sib families were produced with reused males, and were included only in complementary descriptive analyses (Table S2). An average of 52 ± 3 eggs (mean ± standard error, SE) per family were incubated under common garden conditions in darkness, separately in 5 × 5 × 5 cm boxes placed in a 20 L tank. Water temperature in the tank was maintained at 8.1°C ± 0.2°C (mean ± standard deviation) using a water chiller (TK500, TECO, Italy) and was continuously oxygenated using an air pump. This rearing temperature was chosen as a compromise between suboptimal thermal thresholds and experimental stability. It ensured thermal consistency across tanks, maintained high survival rates (84 % Allos and 75 % Geneva), and minimized genotype-by-environment confounding effects by maintaining both populations under identical conditions. Temperature was continuously monitored using a thermal sensor (HOBO UA-002-64, Onset Computer Co., USA). After hatching, alevins were transferred to temperature-controlled tank (8.2°C ± 0.4°C) with continuous aeration using a closed-circuit air pump. Water was filtered, aerated, and partially renewed daily under UV-sterilized conditions. Fish started to feed at approximately 300 ADD (≈37 days post-hatching), and both populations had synchronised developmental timing and therefore started to feed at the same time. Fish were fed *ad libitum* per day with pelletized food (Larviva ProWean 100, BioMar, France).

### 2.4 Morphological traits

We quantified differences in morphology between Lake Allos and Lake Geneva individuals, using both adults (F_0_, n=50) and juveniles (F_1_, 10 individuals per family; Allos, 8 families; Geneva, 6 families; 123-125 days post-hatching). Fish were photographed in lateral view (left side) against a white background using a digital camera (Nikon D5300, Japan). The adults from Allos were euthanized using an excess of benzocaine. To minimize stress and movement during image capture, both adults from Geneva and juveniles were anesthetized using a benzocaine solution prior to photography. We quantified head shape in adults and juveniles using nine homologous landmarks in adults and eight in juveniles, along with twenty-one semi-landmarks (hereafter referred to as landmarks; Figure S2; Table S4; [50]). Landmarks were positioned to capture variation in cephalic morphology [51,52] and digitized using tpsDig2 v.2.32 [53].

### 2.5 Growth

To estimate growth and age of Arctic charr, scales were collected from 27 and 30 adults from lakes Allos and Geneva, respectively. In Lake Allos, individuals correspond to the experimental breeders (excluding one individual with regenerated scales), whereas in Lake Geneva, adult scales were collected independently in 2023 and 2024, as breeders could not be sampled directly. Scales were cleaned, mounted between glass slides, and photographed under transmitted light using a stereomicroscope (Leica S9i, Germany). Interannuli distances were measured on three scales per individuals using ImageJ software (v. 1.54j, NIH, USA; [54]). Only the first four years of growth were retained to ensure consistent comparisons across populations. Since the four annual growth increments (calculated from successive interannuli distances G_0−1_(first year), G_1−2_(second year), G_2−3_(third year) and G_3−4_(fourth year)) were positively correlated, with high associations among the first three years (r = 0.90-0.93, all p < 0.001) and weaker but significant correlations involving the fourth year (r = 0.38-0.54, p ≤ 0.01), they were synthetised in PCA. Only individuals with complete data for all four growth intervals were retained. The first principal component (PC1, 77.8%) was primarily associated with overall growth amplitude, while PC2 (18.4%) captured variation in late growth (G_3−4_; Figure S3).

#### Juvenile specific growth rate (SGR)

Juvenile growth rate was calculated from total length at hatching (alevin stage) to the juvenile stage (123 days post-hatching), after complete yolk-sac resorption and the start of exogenous feeding (300 ADD; ≈ 37 days post-hatching). The total length (Lt, mm) of each individual was measured from the tip of the snout to the end of the caudal fin using ImageJ software. At hatching, 10 individuals per family were randomly collected and photographed in lateral view (Nikon D5300, Sigma 105 mm lens), and the mean family length at hatching was used in subsequent SGR calculations. At the juvenile stage, length was measured individually from the same lateral photographs used for geometric morphometric analyses. Then we quantified specific growth rate (percentage of length increase per unit of time), following [55] as:

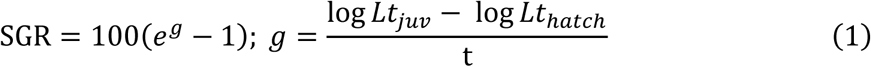

Where 𝐿𝑡_𝑗𝑢𝑣_ is the total length (mm) of juveniles at the time of measurement, 𝐿𝑡_ℎ𝑎𝑡𝑐ℎ_ (mm) is the mean total length at hatching calculated per family, and 𝑡 is the time elapsed between hatching and juvenile measurement in days. To assess potential maternal effects, we quantified yolk sac volume (YSV, mm^3^) as 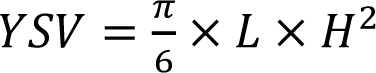, where L and H represent yolk sac major and minor axes, respectively.

### 2.6 Routine metabolic rate

Routine metabolic rate (RMR) was estimated by measuring oxygen consumption rate (ṀO₂) using an intermittent-flow respirometry system [43]. Respirometry measurements were conducted on a minimum of 11 juveniles per family (F1, range: 546-1385 ADD, corresponding to the early parr stage in salmonids), with five families per population randomly selected (Table S4). Measurements were taken at two temperatures (6°C and 10°C), after a one-week acclimation period at those temperatures. These conditions represent, respectively, an ecologically relevant intermediate temperature and a moderately stressful condition within the thermal range experienced by juveniles in their natural habitats (Figure S1). This +4°C increase also reflects the mean temperature rise projected under current climate scenarios [44], and has been used in multiple experimental warming studies (e.g. [45]). Distinct juveniles were used for each temperature.

Fish were fasted for 24 h to eliminate postprandial oxygen consumption [46], weighed to the nearest 0.001 g and introduced into acrylic respirometry chambers (total volume: 55 mL, diameter: 42 mm, height: 40 mm; Loligo Systems, Viborg, Denmark) at the same time every day (11:00 am) for approximately 22 hours. The chambers were placed in the darkness and submerged in two water reservoirs connected to an air pump (Tetra AS150, Germany) to maintain oxygenation, and to a water chiller (TK500, TECO, Ravenna, Italy) to control for temperature. Dissolved Oxygen (DO) concentration (mg·L⁻¹) was recorded every second using a fiber-optic mini-sensor connected to a four-channel oxygen meter (Witrox-4, OX11875, Loligo Systems, Viborg, Denmark), and monitored using AutoResp^TM^ v.3 software (Loligo Systems, Viborg, Denmark). Oxygen consumption rates were calculated based on the slope of oxygen concentration decline over 60 min during a 90 min closed-chamber period. A 3 min flush phase using an automated pump (New Jet NJ800, Newa, Italy) was ensuing water reoxygenation to 100% before each measurement period. The first slope, and those with a coefficient of determination (R^2^) < 0.85 were discarded to account for potential handling stress and to ensure that only stable oxygen consumption rates were analysed [47,48]. The background bacterial oxygen consumption was measured in the absence of fish for one hour before and after each trial in all chambers. The corrected routine metabolic rate (mgO_2_.h^-1^) was obtained by subtracting bacterial respiration using the *convert_rate* function from the R package *respR* (v.2.3.3; [49]).

To minimize potential developmental biases, the tested populations were alternated throughout the experiment over several weeks (Figure S4), and four chambers were placed at each temperature for simultaneous measurements. After respirometry, fish were transferred to separate rearing tanks and were not reused for subsequent phenotypic measurements, to avoid any carry-over or stress effects. The respirometry chambers, tubing and reservoirs were soap-cleaned and dried weekly to prevent the accumulation of bacterial biofilms that could alter measurement accuracy.

### 2.7 Genetic analyses

Fin clips were collected from all adults (n = 50) and preserved in 96% ethanol. DNA was extracted using the QIAwave DNA Blood & Tissue Kit (Qiagen) and the Wizard® SV 96 genomic DNA purification system (Promega), following the manufacturer’s instructions (Appendix S1). A total of 15 neutral microsatellite markers were utilised for this study. Microsatellite genotyping was performed using high-throughput sequencing following the SSRseq approach, as described by Lepais *et al*. (2020) (see Appendix S1 for details). Neutral genetic differentiation was estimated using pairwise-*F*_*st*_based on the method of Weir and Cockerham (1984) implemented in the R package *hierfstat* [56]. Wright’s F-statistics (*F*_*st*_), quantifies the proportion of genetic variance attributable to population structure ranging from 0 (no differentiation) to 1 (complete differentiation) [57]. A 95% confidence interval (CI) for the observed *F*𝑠𝑡 value was estimated using a standard bootstrap procedure with 10,000 iterations.

### 2.8. Statistical analysis

We first aimed at testing phenotypic differences among adult (F_0_) individuals from both populations. To do so, landmark data were analysed using the R package geomorph v. 4.0.10 [58]. Landmark configurations were then aligned using Generalized Procrustes Analysis (GPA) implemented in the function *gpagen* [59]. Centroid size (Csize) was calculated and extracted during GPA and used as a proxy for overall head size. Shape variation was analysed using Procrustes ANCOVA with residual randomization (RRPP; 10,000 permutations) implemented in the function *procD.lm* [60]. Log-transformed centroid size was included as a covariate and population (*pop*) as a fixed factor. Size-only (𝑠ℎ𝑎𝑝𝑒 ∼ log (𝐶𝑠𝑖𝑧𝑒)), reduced (𝑠ℎ𝑎𝑝𝑒 ∼ log (𝐶𝑠𝑖𝑧𝑒) + 𝑝𝑜𝑝) and full models (𝑠ℎ𝑎𝑝𝑒 ∼ log(𝐶𝑠𝑖𝑧𝑒) × 𝑝𝑜𝑝) were compared to test for differences in mean shape and allometric slopes between populations. When the interaction term was not supported, the reduced model was retained. Principal Component Analysis (PCA) was used to visually and synthetised shape variation using *gm.prcomp* implemented in geomorph. PCA scores were used to reduce dimensionality and summarize the major axes of shape variation. These axes were subsequently used as response variables in linear or linear mixed-effects models, and to estimate population differentiation (*P*_*st*_or *Q*_*st*_) relative to neutral genetic differentiation (*F*_*st*_), see below.

We fitted linear models using annual growth PCA axes as dependent variable, and populations (two-levels factor: Allos and Geneva) as an explanatory variable. Then, to determine the juvenile (F_1_) growth (SGR) differences between populations, we used a LMM using the function *lmer* from the *lme4* R-packages [61], with family included as a random effect. We similarly tested whether populations displayed different routine metabolic rate (RMR), metabolic plasticity and body mass scaling using LMM. Before the analysis, RMR (mgO_2_.h^-1^) and body mass (g) were log_10_-transformed to linearize their relationship, as commonly suggested [62–64]. RMR was fitted as a dependent variable, temperature (TP; two-levels factor: 6°C and 10°C), population, body mass (BM) and all possible interactions were included as fixed effect. Family was fitted as a random effect. Non-significant interactions were removed from the model to allow interpretation of lower order interactions and simple terms (Table S5). For all models, residuals were visually checked for normality and homoscedasticity. Significance of fixed effects was assessed using Type-II ANOVA using the *Anova* function from the *car* R-package [65].

### 2.9. Genetic, phenotypic and quantitative differentiation analyses

Finally, we aimed at testing the mechanisms underlying phenotypic differentiation between the populations. To do so, we compared neutral genetic differentiation (*F*_*st*_) to its phenotypic analogues: *Q*_*st*_and *P*_*st*_, which assess differentiation in quantitative traits (i.e., RMR, morphometric traits, growth, SGR) [66,67]. *Q*_*st*_ is based on genetic variance components from a common garden experiment while *P*_*st*_ is as a phenotypic proxy for *Q*_*st*_used on wild animals when no information on parental relatedness are available [68,67]. Briefly, *Q*_*st*_ > *F*_*st*_indicates divergent selection on the considered traits; *Q*_*st*_ < *F*_*st*_ indicates stabilizing selection; and *Q*_*st*_ ≈ *F*_*st*_ indicates differentiation is likely due to genetic drift [67]. The conclusions related to the *P*_*st*_are similar in magnitude, but we cannot tease apart adaptive divergence due to selection and plasticity because of environmental effects.

We estimated *Q*_*st*_ using the following formula:

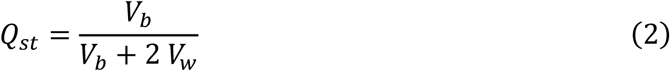

Where 𝑉_𝑏_ represents the between-population genetic variance component and 𝑉_𝑤_ the within population variance component. 𝑉_𝑏_ was estimated as the observed variance component between populations and 𝑉_𝑤_ was approximated by the variance between families (𝑉_𝑓𝑎𝑚_). Given our full-sib design 𝑉_𝑤_ = 2𝑉_𝑓𝑎𝑚_ [69].

All variance components (𝑉_𝑏_, 𝑉_𝑓𝑎𝑚_) were estimated from LMM fitted with population and family (nested within population) as random intercepts (Table S6). To estimate variance components for the relationships between RMR and body mass (BM) or temperature (TP), we fitted separate LMM models with random intercepts and random slopes (BM or TP) at the family level nested within population.

We estimated *P*_*st*_ using the formula proposed by Brommer (2011):

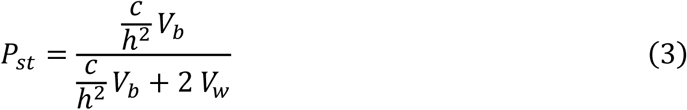

Where 𝑉_𝑏_ and 𝑉_𝑤_ (residual variance) represent the phenotypic variance between and within populations, respectively. 𝑐 corresponds to the proportion of the total variance assumed to be due to additive genetic effects between populations and ℎ^2^ is the heritability. Since we have no information on the c and ℎ^2^ value, we used a conservative 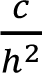ratio of 0.5, and we also performed sensitivity analyses with different values (0.63 and 1; [70]). Both 𝑉_𝑏_ and 𝑉_𝑤_ were extracted from LMM where population was included as a random intercept term.

We assessed whether the observed *P*_*st*_ and *Q*_*st*_ significantly deviated from their expected distribution under neutral hypothesis (see Gilbert & Whitlock, 2015). Briefly, this method uses parametric resampling of genetic variance components (via chi-square distributions) with bootstrapped *F*_*st*_ to simulate the distribution of *Q*_*st*_ (and *P*_*st*_) under a neutral model of genetic drift. For each iteration, the between-population variance under neutrality was calculated using the formula [71]:

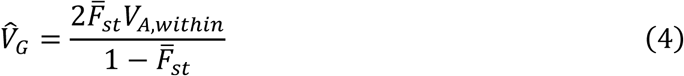

Where 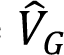 is the variance among populations in the mean value of the trait, 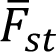 represents the mean *F*_*st*_ estimated across neutral loci and used as a reference for test selection, and 𝑉_𝐴,𝑤𝑖𝑡ℎ𝑖𝑛_ is the additive genetic variance within populations for the trait [71]. We generated 10,000 bootstrap replicates of *Q*_*st*_ under neutrality and constructed the corresponding reference distribution (95% CI). We then calculated the distribution of *Q*_*st*_ − *F*_*st*_ under neutrality differences and its 95% CI. Significance was assessed by evaluating whether the observed difference *Q*_*st*_ − *F*_*st*_ fell outside this neutral distribution, based on empirical *p*-values [71]. A significantly positive difference (p < 0.05) was interpreted as evidence of divergent selection, whereas a significantly negative difference indicated stabilizing selection [71].

All statistical analyses were performed using R language (v. 4.4.2; [72]).

## 3. Results

### 3.1. Morphological differentiation

Adult morphology differed markedly between populations (R^2^ = 0.36, F = 27.47, p < 0.001; Table 1, Figure 2). Head size (centroid size) did not significantly explain head shape variation (R^2^ = 0.02, F = 1.78, p = 0.11), and the interaction between size and population was not supported (R^2^ = 0.008, F = 0.60, p = 0.72). Shape differences were primarily expressed along PC1 (45.4%), which described variation in head size (Figure 2A). Specifically, individuals from Lake Geneva exhibited more compact and deeper head shapes with a relatively reduced anterior projection, whereas fish from Lake Allos showed more elongated head shapes with a slightly extended rostral region, as indicated by higher values in PC1 (Figure 2A). Morphology synthetised by PC2 (19.4%) captured small variation in dorsal curvature and orbital region position but did not structure the separation between populations (Figure 2A).

**Table 1.**
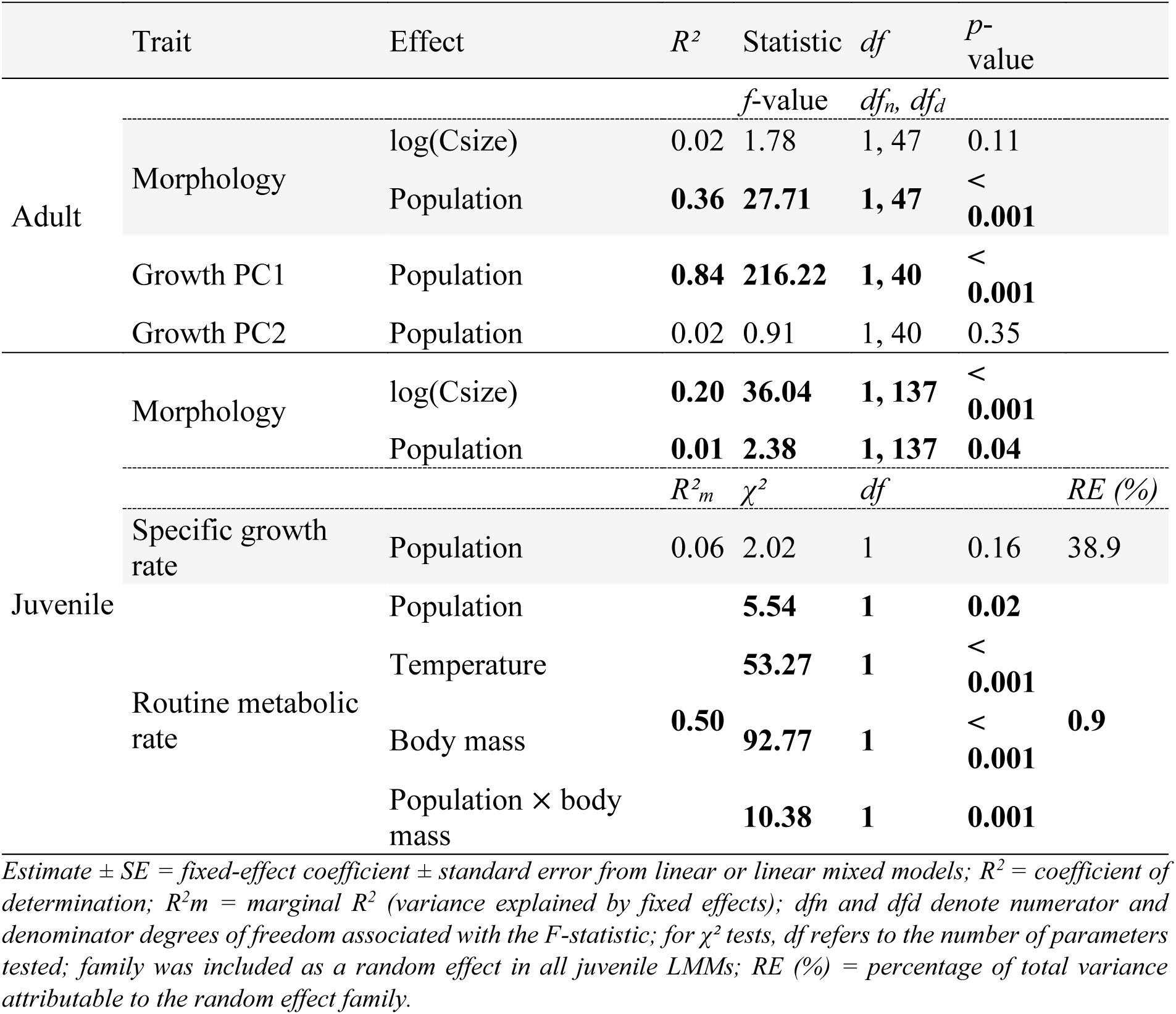
Effects of population of origin on morphology and growth, and effects of temperature treatments (6°C and 10°C), population (Allos; Geneva) and body mass (g, log10-transformed) and their interaction on juvenile routine metabolic rate (RMR; mgO2.h-1, log10-transformed) and specific growth rate (SGR) in Arctic charr (*Salvelinus alpinus*). Adult and juvenile shape were analysed using Procrustes ANCOVA. Adult growth was analysed using linear models. Juvenile RMR and SGR were analysed using linear mixed-effects models with family included as a random effect. Significance is indicated in bold (*p* < 0.05).

**Figure 2.**
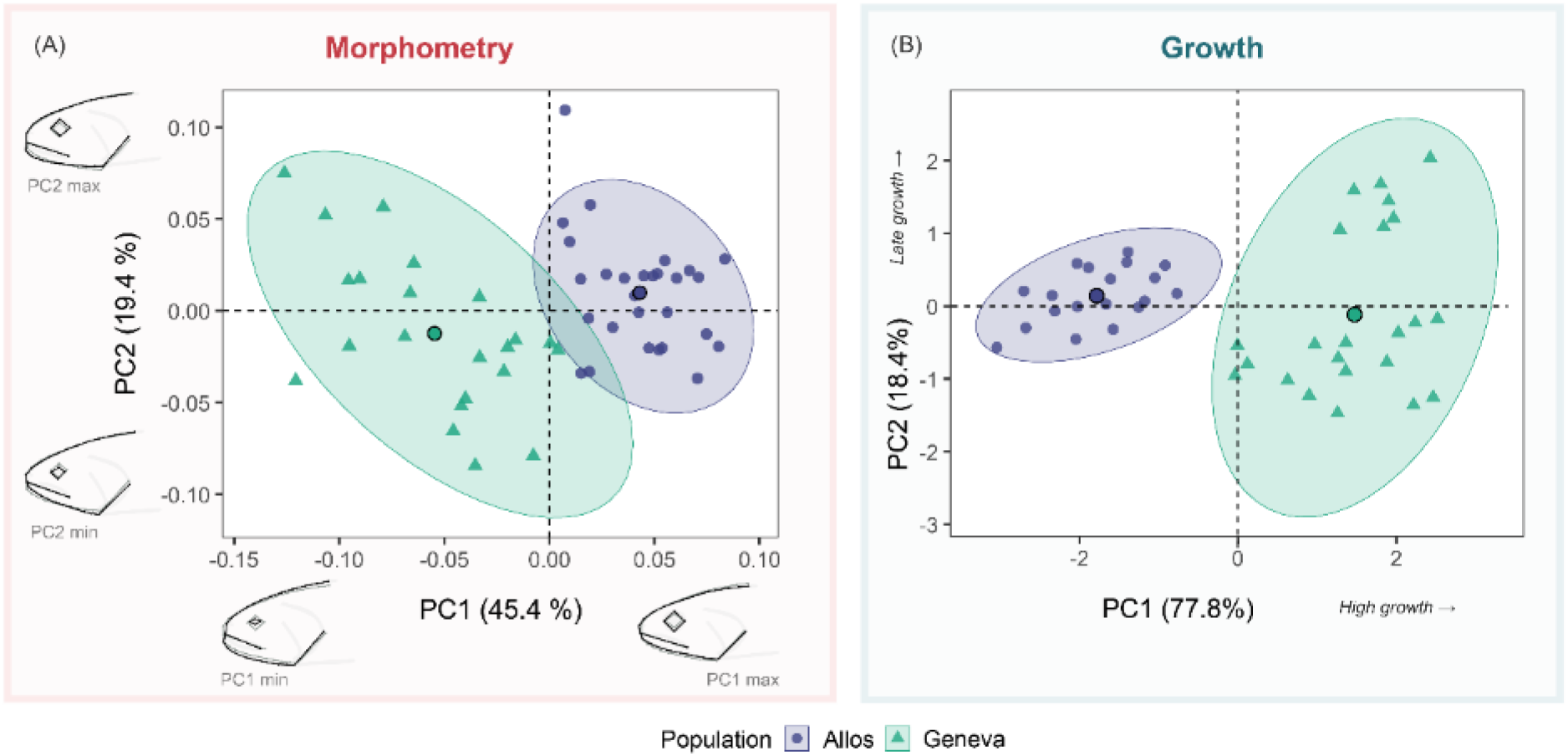
Differentiation in shape and growth of adult Arctic charr (*Salvelinus alpinus*) from Lakes Allos and Geneva. (A) Principal component analysis (PCA) of size-corrected Procrustes shape coordinates. Thin-plate spline deformation grids illustrate shape variation along PC1 and PC2. Deformations are shown relative to the reference configuration and magnified two-fold for clarity. Grey outlines represent the reference (mean) shape, and black lines represent the two extremes along each principal component axis. (B) PCA based on four annual growth increments estimated from scale radius. Each point represents an individual (Allos, blue circles; Geneva, green triangles). Ellipses indicate 95% confidence intervals and larger points represent the multivariate mean (centroid) of each population. Arrows indicate the direction of increasing growth.

In juveniles reared under common garden conditions, shape showed a small but significant difference between populations (R^2^ = 0.01, F = 2.38, p < 0.04; Figure 3A, Table 1). In contrast, centroid size explained a substantially larger proportion of shape variation (R^2^ = 0.20, F = 36.04, p < 0.001), and the interaction between size and population was not supported (R^2^ = 0.01, F = 1.25, p = 0.25).

**Figure 3.**
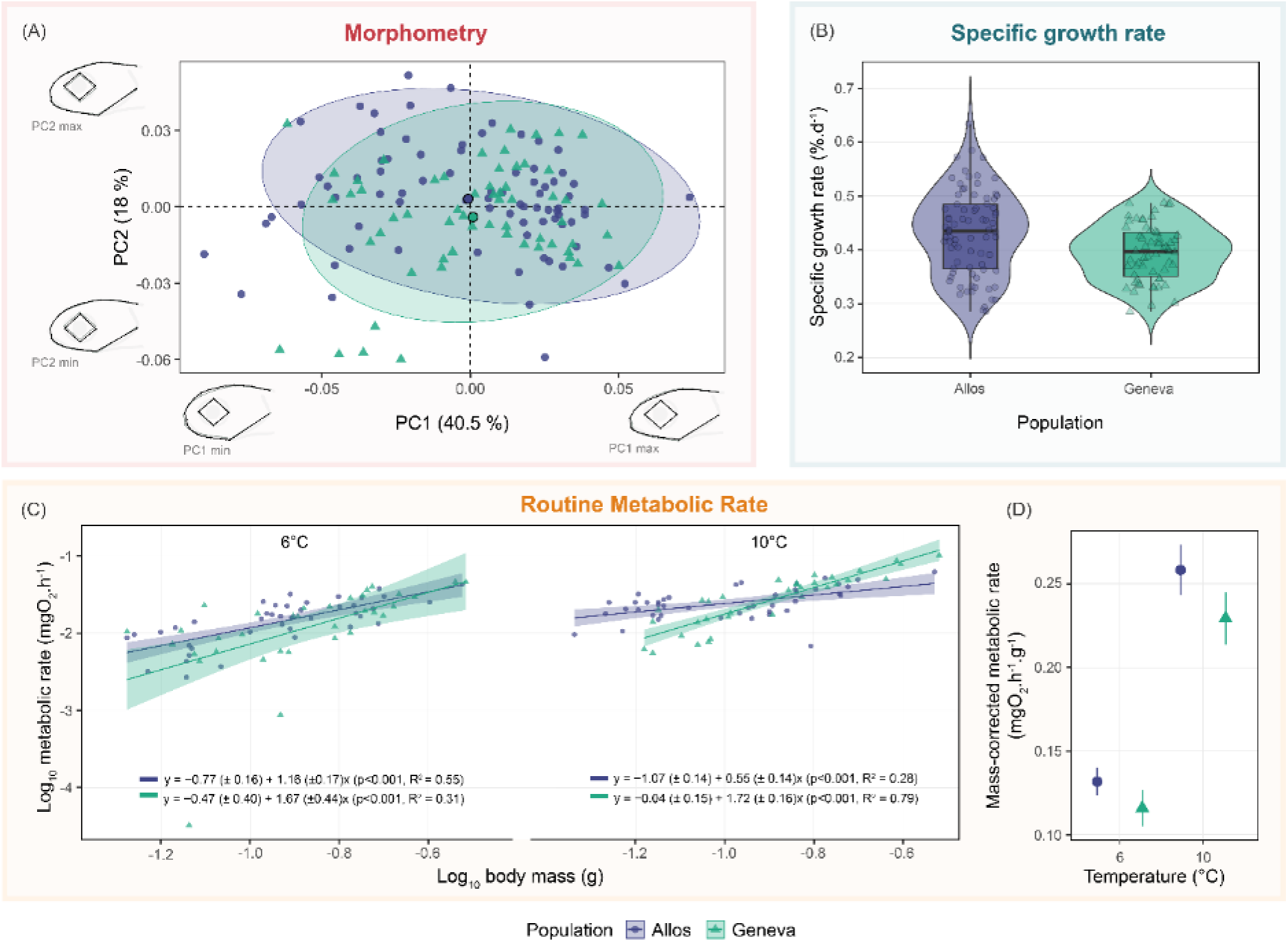
Multi-traits comparison of juvenile Arctic charr (*Salvelinus alpinus*) from Lakes Allos and Geneva under common garden conditions. (A) Principal component analysis (PCA) of size-corrected Procrustes shape coordinates. Thin-plate spline deformation illustrate shape variation along PC1 and PC2. Deformations are shown relative to the mean shape (grey outline) and black lines represent extremes along each axis, magnified two-fold for clarity. (B) Specific growth rate (SGR, %.d^-1^) measured from hatching to the juvenile stage. Violin plots depict the population distribution (Allos, n = 80; 8 families; Geneva, n = 60; 6 families). Axes are annotated with the main contributing traits (arrows indicate direction of increase). (C) Relationship between log_10_ -transformed routine metabolic rate (mgO_2_ .h^-1^) and log_10_ -transformed body mass (g) at 6°C and 10°C for each population. Scaling exponents (± SE), *p*-values and R² are indicated by population and temperature. Each point represents an individual (Allos, blue circles; Geneva, green triangles), lines and shaded indicate fitted linear regressions with 95% confidence intervals. (D) Mean mass-corrected routine metabolic rate (mgO_2_ .h^-1^.g^-1^) by population and temperature with error bars indicating standard errors (± SE).

### 3.2. Growth

Adults from Lake Geneva displayed higher growth rates than adults from Lake Allos, especially during the first three years of life of individuals (growth axis 1, R^2^ = 0.84, F = 216.22, p < 0.001; Table 1, Figure 2B). Growth axis 2, associated with late growth variation, did not differ significantly between lakes (Table 1).

Juveniles from both populations displayed similar specific growth rates (% per day; mean ± SE = 0.432 ± 0.017 % in Allos and 0.395 ± 0.020 % in Lake Geneva; χ^2^ = 2.02, p = 0.16; Figure 3B). Similarly, yolk sac volume measured at hatching did not differ significantly between populations (64 ± 3.48 mm^3^ in Allos and 66.3 ± 4.01 mm^3^ in Lake Geneva; χ^2^ = 0.20, p = 0.65; Figure S5).

### 3.3. Routine metabolic rate

Routine metabolic rate differed between populations (χ^2^ = 5.5, p = 0.019; Figure 3C, D), with Allos individuals consistently showing higher mass-corrected rates (mean RMR ± SE = 0.258 ± 0.015) than Geneva individuals (0.229 ± 0.016; Figure 3D). Moreover, juveniles from Lake Geneva displayed a higher allometric exponent between RMR and body mass than juveniles from Lake Allos (significant interaction between population and body mass; χ^2^ = 10.4, p = 0.001; Figure 3C). At 6 °C, the scaling exponent was 1.16 ± 0.17 in Lake Allos and 1.67 ± 0.44 in Lake Geneva, whereas at 10 °C it was 0.55 ± 0.14 in Lake Allos and 1.72 ± 0.16 in Lake Geneva (Figure 3C). Finally, the thermal reaction norm of metabolic rate was similar between the populations, as indicated by the non-significant interaction between temperature and populations (χ^2^ = 1.30, p = 0.25; Table S5). Overall, the model explained about 50% of the total variation in RMR (R^2^m = 0.50).

### 3.4. Genetic and quantitative differentiation

Neutral genetic differentiation between the two populations was low (*F*_*st*_ = 0.017; 95% CI: 0.006-0.033; Table S7).

In adults, phenotypic differentiation among populations relative to morphometric PC1 (*P*_*st*_ = 0.45) was significantly higher than neutral genetic differentiation (p < 0.001; Figure 4, Table S6). This result was similar under conservative assumptions about trait heritability (*P*_*st*_ = 0.55 when c/h² = 0.5) and under low heritability (*P*_*st*_ = 0.71 when c/h² = 1). In contrast, population differentiation along morphometric PC2 did not differ from expectations of genetic drift only (*P*_*st*_ = 0.02, Figure 4, Table S6). Annual growth trajectories differed significantly between populations, with a high *P*_*st*_ growth axis 1 (*P*_*st*_ = 0.63, p < 0.001), but not in late-life growth (axis 2, Table S6).

**Figure 4.**
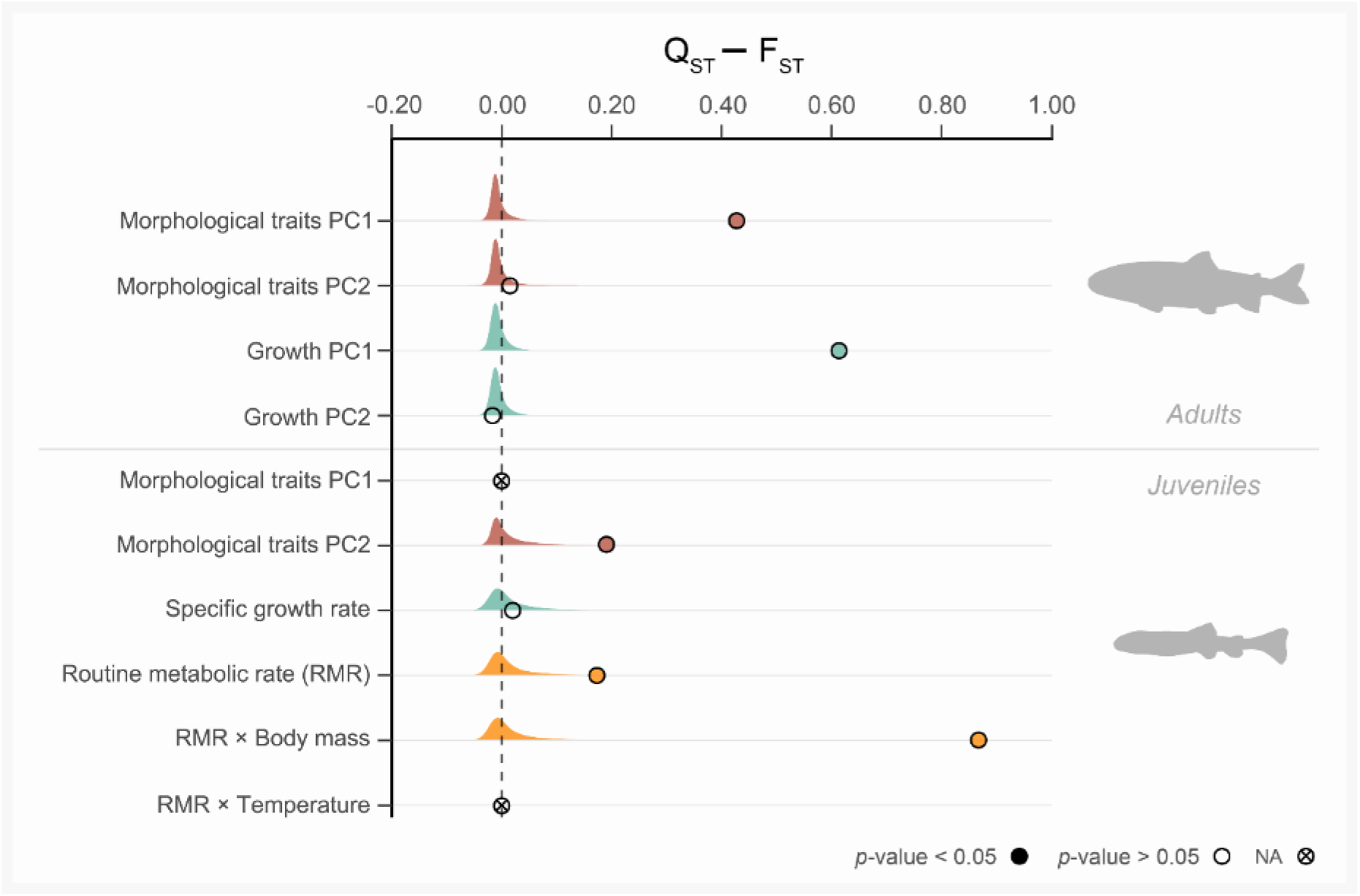
Simulated neutral distributions and observed values of *P*_*st*_ − *F*_*st*_ for 2 traits in adults and of *Q*_*st*_ − *F*_*st*_ for 5 traits in juveniles Arctic charr (*Salvelinus alpinus*) from Lake Allos and Geneva. The density curves represent the expected distribution of *P*_*st*_ − *F*_*st*_ and *Q*_*st*_ − *F*_*st*_ under neutrality, simulated following Whitlock and Guillaume (2009), based on empirical estimates of *F*_*st*_ and within-population variance for each trait. Observed values are shown as points: solid circles indicate significant deviations from neutrality (*p* < 0.05), open circles indicate non-significant deviation and crossed circles correspond to traits for which the test was not applicable due to the absence of between-population variance (PC1 in juvenile morphology). The vertical dashed line indicates the neutral expectation *P*_*st*_ − *F*_*st*_ and *Q*_*st*_ − *F*_*st*_. Traits are grouped by functional category (morphology: red; growth: green, metabolism: orange) and separated between juveniles and adults.

Population differentiation in quantitative traits varied across traits (Figure 4). Juvenile shape showed moderate differentiation along PC2 (*Q*_*st*_ = 0.21, p = 0.01; Table S6). In contrast, no *Q*_*st*_ estimate was calculated for PC1 because the linear mixed-effects model showed no detectable among-population variance, with all variances attributed to the residual component. Routine metabolic rate showed significant differentiation between populations (*Q*_*st*_ = 0.19, p = 0.01; Figure 4, Table S6). Similarly, the differentiation in the scaling exponent was significantly higher than *F*_*st*_ (*Q*_*st*_ = 0.88, p < 0.001), while no differentiation was detected for the thermal reaction norm. Differentiation in SGR (*Q*_*st*_ = 0.04, p = 0.26) did not significantly differ from *F*_*st*_ (Table S6).

## 4. Discussion

Rapid phenotypic divergence under recent allopatric isolation offers valuable opportunities to study how plasticity and selection interact to drive early stages of adaptation [7,28]. By combining field, experimental and quantitative genetic approaches, we found that two Arctic charr populations, isolated for approximately a century and exposed to contrasting environments, exhibit phenotypic divergence that exceeds expectations under neutral genetic drift alone. These findings indicate that phenotypic differentiation in key traits related to morphology, metabolism and growth resulted from both plasticity and adaptive selection.

Our findings suggest that population divergence in morphological traits was mainly driven by plasticity. Indeed, while adults strongly differed in their morphology (i.e., supported by significant *P*_*st*_ values), the common garden experiment indicated only weak morphological differentiation among juveniles, with extensive overlap in head shape between populations (Figure 4C), suggesting that these traits were poorly associated with genetic components [71]. This pattern suggests phenotypic plasticity shaped by the contrasted environments of the lakes. Moreover, growth trajectories between populations were highly divergent. Individuals from Lake Geneva grew faster than individuals from Lake Allos, reflecting contrasting environmental effects potentially linked with local resource availability [33]. In high-altitude Alpine lakes, growth may be constrained not by maximal summer temperatures but rather by the short post ice-off window of prey availability (typically following 5-6 months of ice cover at Lake Allos), emphasizing food limitation as the primary constraint on somatic development in low-productivity environments [73].

Interestingly, we showed physiological divergences between juveniles from different populations. Individuals from Allos exhibited higher RMR, and a lower allometric exponent than those from Geneva. These patterns were consistent across tested temperatures (6°C and 10°C). This may reflect physiological constraints at warmer temperatures in a population adapted to a colder environment, given that normal natal thermal conditions are generally colder in Lake Allos, (i.e., 4-5°C see Figure S1). Moreover, populations displayed different scaling exponents between RMR and body mass. The interaction of temperature and body mass on metabolism has been shown in multiple species and we here demonstrated that it can evolve rapidly between populations. This probably results from the selection of different reaction-norms of scaling exponent among populations, implying size-dependent metabolic costs [74]. Cold-adapted fish populations may show reduced metabolic scaling exponents at higher temperatures, particularly at higher body mass, potentially due to limitations in tissue oxygenation or energy-conservative strategies favouring efficiency over performance [74]. Other factors, such as resource supply or lifestyle, are important in shaping scaling exponent and further studies may integrate multiple environmental characteristics to better understand evolution of scaling exponent [74]. The apparent divergence between juvenile physiological responses and adult growth may reflect environmental constraints limiting realized growth in the wild. In colder and low-productivity environments such as Lake Allos [37], individuals are expected to develop physiological strategies improving energy-use efficiency [75]. When constraints are relaxed under common conditions with abundant resources, intrinsic differences between populations may become more apparent, consistent with countergradient variation [76].

Such size-dependent energetic requirements may provide a mechanistic link between metabolic divergence and the observed growth trajectories. Especially, at 10°C, this higher metabolic rate and attenuated metabolic scaling in Allos individuals was associated with a slightly higher SGR, suggesting accelerated development. Although initially showing little difference, SGR significantly differed between populations when a broader sample was considered (see Figure S6). The coupling of accelerated early growth and reduced metabolic scaling aligns with the temperature-size rule (TSR; [77]), which predicts enhanced growth responses under warmer-than-natal conditions. It also aligns with the predictions of multiphasic ontogenetic allometry [64], which posits that the transition from cellular proliferation to differentiation is accompanied by a decrease in the allometric slope of metabolism. Incorporating multigenerational approaches with controlled crosses across multiple generations would allow more robust inference on these heritable mechanisms [4].

Physiological divergence was further supported by *Q*_*st*_ − *F*_*st*_ comparisons, suggesting selection on metabolic traits, including components associated with body mass. This pattern may be attributed to transgenerational developmental plasticity, parental effects or directional selection, as all measurements were conducted on F_1_ generation [78]. While maternal effects may have contributed to phenotypic differences, the similar yolk sac volume (maternal provisioning trait) between populations (Figure S5) suggested that this effect was limited.

In addition, transgenerational plasticity may be involved, whereby parental environmental conditions induce phenotypic changes in the offspring, such as in brook charr (*Salvelinus fontinalis*), transcriptional responses in fry have been shown to reflect the thermal history of their parents, despite a limited contribution of additive genetic variance [79]. Together, our results suggest that early-life metabolic differences may represent key targets of selection under thermal divergence, potentially triggering cascading effects on growth and ontogenetic development. Incorporating both ontogenetic and transgenerational perspectives may provide further insights regarding the physiological adaptation to climate change.

All these processes diverged in less than a century through the combined effect of plasticity and selection. These findings allow a better understanding of the evolution of marginal populations (i.e., lake Allos), which may either foster rapid adaptation or face constraints due to ecological stress (e.g., extreme conditions, isolation) [33,34]. However, the Allos population showed no evidence of reduced genetic diversity and was genetically similar to the Geneva population (see Table S7). Lake Allos is characterized by geographic isolation, low productivity and cold temperatures, as commonly observed in mountain lakes [80]. This population displays physiological and morphological differentiation, highlighting high adaptive potential under environmental constraints. While often demographically disadvantaged, peripheral habitats may play a crucial role in adaptive diversification by enabling strong selection on key traits for local persistence [81]. This resonates with findings in other marginal salmonid populations, reinforcing the notion that edge-of-range environments can act as sentinels for rapid adaptation in the face of climate-driven pressures [32]. While genetic differentiation based on neutral markers was low, further investigating candidate genes might generate mechanistic insights regarding the evolutionary processes driving of trait divergence [82].

In conclusion, this study highlights the complex interplay between phenotypic plasticity and natural selection in contexts of recent allopatric isolation. Our findings emphasise the value of integrative functional approaches linking metabolic, growth, and morphological traits to better understand how populations respond to novel thermal environments. Despite the short timescales involved, we detected significant divergence in physiological traits, suggesting that early-life metabolism can evolve rapidly under divergent selection pressures. This supports the idea that evolutionary responses to environmental variability may occur over contemporary timescales, particularly in edge-of-range populations where selection and genetic drift interact and may provide early signals of adaptive change under global stressors.

## Acknowledgements

The study was carried out in an experimental technical facility provided by the Observatory of LAkes (OLA) part of AnaEE-France (Analysis and Experimentation on Ecosystems) and LIFE Research Infrastructure (IR LIFE, INRAE). We thank the Rive fish farm for providing gametes from Lake Geneva individuals and to the Mercantour National Park for granting fishing permissions in Lake Allos. We are particularly grateful to Valérie Hamelet for her invaluable assistance with scalimetry analyses and to Cécile Chardon for her support with DNA extractions. We also thank Andréa-Marie Lambert for her help with the setup of metabolic experiments. Finally, we also thank Christophe Boury, Olivier Lepais and Erwan Guichoux for their contribution to the development of microsatellite markers, and Jérôme Prunier for his support with bioinformatic processing, as well as Matthias Vignon for his assistance with geometric morphometrics. Finally, we thank the four anonymous reviewers for their constructive comments and suggestions, which improved earlier versions of the manuscript.

## Author contributions

Hervé Rogissart: Conceptualization; Methodology; Data curation; Formal analysis; Investigation; Visualization; Writing-original draft; Writing-review & editing. Emilie Chancerel: Methodology (development of microsatellite markers), Writing-review & editing. François-Raphael Lubin: Methodology (support with fish sampling, rearing and metabolic rate measurements). Martin Daufresne: Funding acquisition, Conceptualization, Writing-review & editing. Guillaume Evanno: Methodology (advice on genetic quantitative methods); Writing-review & editing. Jean Guillard: Funding acquisition; Conceptualization, Supervision; Writing-review & editing. Allan Raffard: Funding acquisition; Project administration; Supervision; Conceptualization; Methodology; Investigation; Writing-review & editing.

## Data accessibility

Data will be available on Figshare upon acceptance of the manuscript.

