## Supplementary Material and Appendix for "A century of allopatry: plasticity and rapid selection shape phenotypic trait variability under contrasting environments"

Table S1. Environmental characteristics of Lakes Allos and Geneva. Elevation is given in meters above sea level (m a.s.l.). Mean water temperature corresponds to the natural conditions recorded at spawning depths (10-20 m and 40-60 m for Allos and Geneva, respectively) over the theoretical spawning and incubation period (November to January and December to March, respectively; see Fig. S1).

| **Parameter** | **Allos** | **Geneva** |
|---|---|---|
| Latitude (°N) | 44°14'02" | 46°26'37" |
| Longitude (°E) | 6°42'29" | 6°31'15" |
| Elevation (m a.s.l.) | 2232 | 372 |
| Surface area (km$^2$) | 0.54 | 580 |
| Max. depth (m) | 51 | 309 |
| Mean depth (m) | 25 | 153 |
| Mean water temperature – theoretical spawning period (°C ± SD) | 4.22 ± 1.17 | 7.89 ± 0.68 |
| Mean water temperature theoretical incubation period (°C ± SD) | 3.60 ± 0.16 | 7.34 ± 0.67 |
| Species | TRF, OBL, CRI, VAI | TRF, OBL, VAI, COR, PER, GAR, BRO* |

Note: TRF, *Salmo trutta*; OBL, *Salvelinus alpinus*; CRI, *Salvelinus namaycush*; VAI, *Phoxinus phoxinus*; COR, *Coregonus spp*.; PER, *Perca fluviatilis*; GAR, *Rutilus rutilus*; BRO, *Esox lucius*. Species composition for Lake Allos follows data provided by the regional fisheries federation [1]. For an exhaustive list of species present in Lake Geneva please refer to [2].

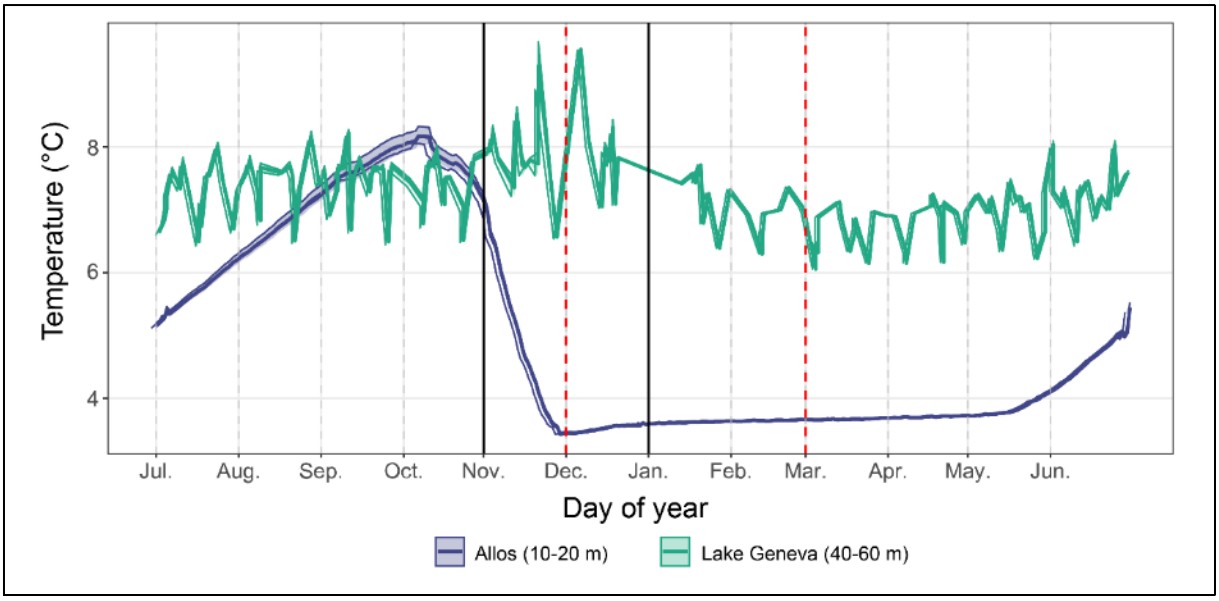


Figure S1. Daily mean water temperature profiles (°C) for Lake Allos (10–20 m depth, blue) and Lake Geneva (40–60 m depth, green), averaged over 2016-2022. Solid lines represent average daily temperatures, and shaded areas indicate the 95% confidence interval. Solid black vertical lines indicate the estimated spawning period (November to early January), and red dashed lines indicate the estimated incubation period of Arctic charr (*Salvelinus alpinus*) eggs, which typically occurs between early December to early March. During the spawning period, mean temperature was 4.22 ± 1.17 °C (SD) in Lake Allos and 7.89 ± 0.68 °C (SD) in Lake Geneva, while during the incubation period it was 3.60 ± 0.16 °C (SD) and 7.34 ± 0.67 °C (SD), respectively. Data were collected for lake Allos from Réseau National de suivi de la Température des plans d'eau (RNT plans d'eau), coordinated by pole R&D ECLA (OFB–INRAE–USMB) and for Lake Geneva from OLA-IS [3], AnaEE-France, INRAE, Thonon-les-Bains and CIPEL.

Table S2. Cross-design scheme to produce F1 offsprings of Arctic charr (*Salvelinus alpinus*) from wild-caught spawners collected using gillnets in two populations (Allos and Geneva). All families were reared under identical temperature conditions (8°C). Families highlighted in grey represent the F1 juveniles used in the Routine Metabolic Rate (RMR) experiment (see Appendix S1: Table S1 for details). Only independent full-sib families were used in quantitative analyses; half-sib families resulting from reused males were included only for complementary descriptive purposes (see also Fig. S6).

| Population | Female | Total length (mm) | Male cross | Length (mm) | Design | Family |
|---|---|---|---|---|---|---|
| Allos | ALO1 | 225 | ALO11 [a] | 267 | Full-sib | A1 |
| Allos | ALO2 | 257 | ALO12 [b] | 250 | Full-sib | A2 |
| Allos | ALO3 | 274 | ALO13 | 239 | Full-sib | A3 |
| Allos | ALO4 | 235 | ALO14 | 241 | Full-sib | A4 |
| Allos | ALO5 | 240 | ALO15 | 264 | Full-sib | A5 |
| Allos | ALO6 | 234 | ALO16 | 221 | Full-sib | A6 |
| Allos | ALO7 | 245 | ALO17 | 284 | Full-sib | A7 |
| Allos | ALO8 | 245 | ALO18 | 250 | Full-sib | A8 |
| Allos | ALO9 | 230 | ALO12 [b] | 250 | Half-sib | A9 |
| Allos | ALO10 | 234 | ALO11 [a] | 267 | Half-sib | A10 |
| Geneva | LEM1 | 520 | LEM10 [c] | 405 | Full-sib | L1 |
| Geneva | LEM2 | 485 | LEM11 [d] | 420 | Full-sib | L2 |
| Geneva | LEM3 | 473 | LEM12 | 430 | Full-sib | L3 |
| Geneva | LEM4 | 475 | LEM13 | 351 | Full-sib | L4 |
| Geneva | LEM5 | 438 | LEM14 [e] | 394 | Full-sib | L5 |
| Geneva | LEM6 | 453 | LEM15 | 421 | Full-sib | L6 |
| Geneva | LEM7 | 491 | LEM14 [e] | 394 | Half-sib | L7 |
| Geneva | LEM8 | 437 | LEM10 [c] | 405 | Half-sib | L8 |
| Geneva | LEM9 | 430 | LEM11 [d] | 420 | Half-sib | L9 |
| Allos | 10 | - | 10 | - | - | 10 |
| Geneva | 9 | - | 9 | - | - | 9 |

[a-e] indicate repeated use of the same male individual across different crosses.

Table S3. Hatching date and degree-days at hatching for F1 offspring of Arctic charr (*Salvelinus alpinus*) from the Allos and Geneva populations. Offspring were incubated under identical laboratory conditions.

| Population | Hatching date | Degree-days at hatch | Family |
|---|---|---|---|
| Allos | 19/01/2024 | 454 | A1 |
| Allos | 18/01/2024 | 454 | A2 |
| Allos | 18/01/2024 | 454 | A3 |
| Allos | 19/01/2024 | 454 | A4 |
| Allos | 19/01/2024 | 454 | A5 |
| Allos | 19/01/2024 | 454 | A6 |
| Allos | 18/01/2024 | 454 | A7 |
| Allos | 19/01/2024 | 454 | A8 |
| Geneva | 18/01/2024 | 453 | L1 |
| Geneva | 17/01/2024 | 439 | L2 |
| Geneva | 18/01/2024 | 449 | L3 |
| Geneva | 19/01/2024 | 455 | L4 |
| Geneva | 19/01/2024 | 458 | L5 |
| Geneva | 19/01/2024 | 455 | L6 |

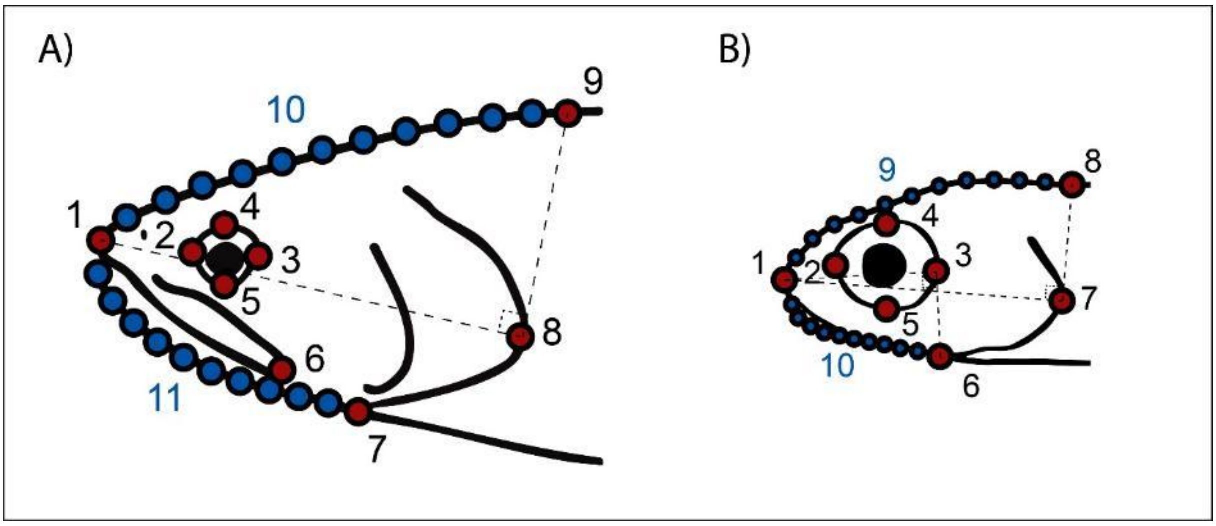


Figure S2. Landmarks (blue) and semi-landmarks (red) used to characterise head shape variation in Arctic charr from Allos and Geneva populations. Configurations are shown for (A) adults and (B) juveniles. For detailed descriptions, refer to Table S4.

Table S4. List of the morphological landmarks (LM) and semi-landmarks (SLM) used in the present study for adults and juveniles.

| Stage | ID | Details on placement |
|---|---|---|
| Adult | 1 | Anterior tip of upper jaw (premaxilla) |
| | 2 | Anterior margin of orbit (horizontal axis) |
| | 3 | Posterior margin of orbit (horizontal axis) |
| | 4 | Dorsal margin of orbit (vertical axis) |
| | 5 | Ventral margin of orbit (vertical axis) |
| | 6 | Posterior edge of maxilla |
| | 7 | Intersection between the ventral edge of the operculum and the ventral body outline |
| | 8 | Anterior margin of operculum |
| | 9 | Intersection between the dorsal body outline and the perpendicular projection from LM1 to LM8 |
| | 10 | Evenly spaced points along the dorsal head and body outline (SLM) |
| | 11 | Evenly spaced points along the ventral head and body outline (SLM) |
| Juvenile | 1 | Anterior tip of upper jaw (premaxilla) |
| | 2 | Anterior margin of orbit (horizontal axis) |
| | 3 | Posterior margin of orbit (horizontal axis) |
| | 4 | Dorsal margin of orbit (vertical axis) |
| | 5 | Ventral margin of orbit (vertical axis) |
| | 6 | Intersection between the ventral body outline and the perpendicular projection from LM1 to LM3 |
| | 7 | Anterior margin of operculum |
| | 8 | Intersection between the dorsal body outline and the perpendicular projection from LM1 to LM7 |
| | 9 | Evenly spaced points along the dorsal head and body outline (SLM) |
| | 10 | Evenly spaced points along the ventral head and body outline (SLM) |

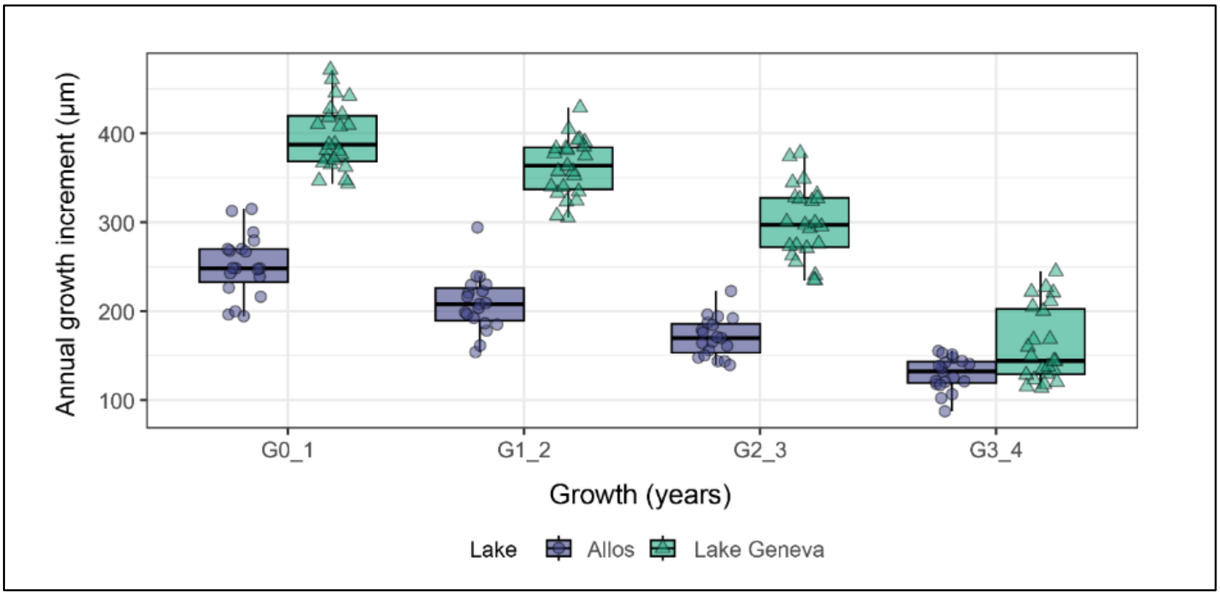


Figure S3. (A) Annual growth increments estimated from scale radius for each growth interval ($G_{0-1}$ to $G_{3-4}$) in Allos (blue) and Lake Geneva (green). Each point corresponds to an individual fish.

Table S4. Summary of the experimental design used in the Routine Metabolic Rate (RMR) experiment on juvenile Arctic charr (*Salvelinus alpinus*). The table reports the families tested in each population (Allos and Geneva) and the number of F1 juveniles measured at each temperature (6 °C and 10 °C).

| Population | Families | Number of juveniles (6°C) | Number of juveniles (10°C) |
|---|---|---|---|
| Allos | A1 | 8 | 8 |
| | A2 | 8 | 8 |
| | A3 | 12 | 11 |
| | A4 | 8 | 8 |
| | A5 | 5 | 8 |
| Geneva | L1 | 7 | 6 |
| | L2 | 8 | 7 |
| | L3 | 8 | 8 |
| | L4 | 6 | 6 |
| | L5 | 5 | 6 |
| **Total (Allos)** | **5** | **41** | **43** |
| **Total (Geneva)** | **5** | **34** | **33** |

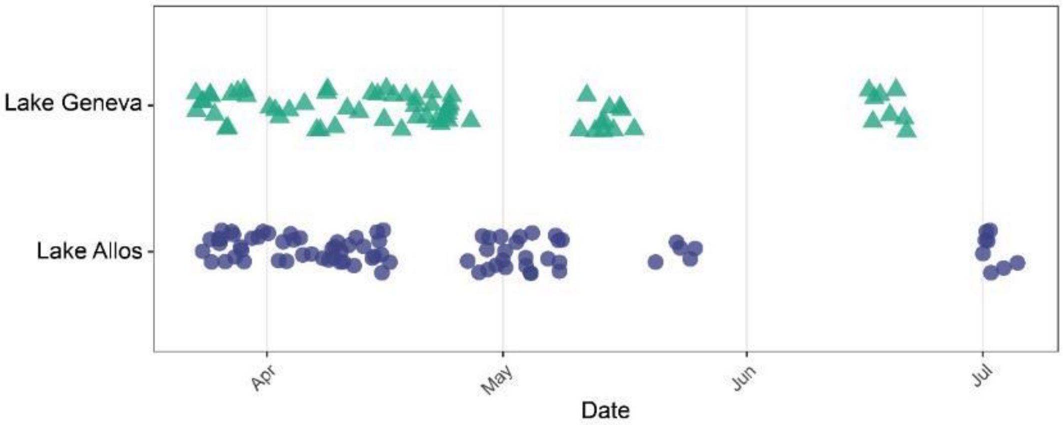


Figure S4. Chronology of respirometry measurements in the two populations. Each point represents an individual measurement day for either Allos (blue circles) or Lake Geneva (green triangles). Populations were tested alternately throughout the experiment, ensuring comparable developmental stages. Measurement dates did not differ between populations ($t = 0.02$, $df = 146$, $p = 0.99$). To further assess whether developmental time influenced metabolic rate, accumulated degree days (ADD) were included as a covariate in the linear mixed-effects model. ADD did not explain variation in metabolic rate (estimate = 0.0048 ± 0.032 SE; $\chi^2 = 0.02$, $p = 0.88$), and inclusion of ADD did not alter the magnitude or significance of temperature (estimate = 0.336 without ADD vs. 0.335 with ADD), population (estimate = 0.638 vs. 0.636), or population x body mass effects (estimate = 0.819 vs. 0.816). The more parsimonious model without ADD was therefore retained.

Table S5. Comparison of linear mixed-effects model (LMMs) explaining variation in routine metabolic rate (RMR) with the family as random effect. Models include different combinations of population (Allos vs. Geneva), temperature treatments (6°C vs. 10°C) and body mass (unit, $log_{10}$-transformed) and their interaction on routine metabolic rate (unit, $log_{10}$-transformed) in Arctic charr juveniles. Model 5 additionally included accumulated degree days (ADD) as a continuous covariate to test for potential developmental effects. Significant is indicated in bold ($p < 0.05$). The most supported model (Model 1) was selected based on the lowest Akaike Information Criterion (AIC = 72.31).

| **Model** | **AIC** | ***df*** | $R^2m$ | $\chi^2$ | ***p*-value** | ***RE (%)*** |
|---|---|---|---|---|---|---|
| **Model 1** | **72.31** | **7** | **0.51** | | | **0.9** |
| population | | 1 | | 5.54 | **0.019** | |
| temperature | | 1 | | 53.27 | **< 0.001** | |
| body mass | | 1 | | 92.77 | **< 0.001** | |
| population × body mass | | 1 | | 10.38 | **0.001** | |
| Model 2 | 75.94 | 8 | 0.51 | | | 0.8 |
| population | | 1 | | 5.59 | **0.02** | |
| temperature | | 1 | | 53.35 | **< 0.001** | |
| body mass | | 1 | | 92.25 | **< 0.001** | |
| population × body mass | | 1 | | 11.04 | **< 0.001** | |
| population × temperature | | 1 | | 1.30 | 0.25 | |
| Model 3 | 77.07 | 9 | 0.52 | | | 1.5 |
| population | | 1 | | 4.99 | **0.03** | |
| temperature | | 1 | | 53.94 | **< 0.001** | |
| body mass | | 1 | | 89.47 | **< 0.001** | |
| population × body mass | | 1 | | 9.88 | **0.002** | |
| population × temperature | | 1 | | 1.84 | 0.17 | |
| temperature × body mass | | 1 | | 1.86 | 0.17 | |
| Model 4 | 76.92 | 10 | 0.52 | | | 1.1 |
| population | | 1 | | 5.20 | **0.02** | |
| temperature | | 1 | | 54.04 | **< 0.001** | |
| body mass | | 1 | | 91.88 | **< 0.001** | |
| population × body mass | | 1 | | 10.25 | **0.001** | |
| population × temperature | | 1 | | 1.85 | 0.17 | |
| temperature × body mass | | 1 | | 1.84 | 0.17 | |
| population × temperature × body mass | | 1 | | 1.73 | 0.19 | |
| Model 5 | 79.31 | 8 | 0.50 | | | 1.0 |
| population | | 1 | | 5.12 | **0.02** | |
| temperature | | 1 | | 51.54 | **< 0.001** | |
| body mass | | 1 | | 45.93 | **< 0.001** | |
| population × body mass | | 1 | | 10.12 | **0.001** | |
| ADD | | 1 | | 0.02 | 0.883 | |

*$R^2m$ = marginal $R^2$ (variance explained by fixed effects); for $\chi^2$ tests, degree of freedom (df) refer to the number of parameters tested; family was included as a random effect in all juvenile LMMs; RE (%) = percentage of total variance attributable to the random effect family.*

Table S6. Summary of the linear mixed models (LMMs) assessing the variance components and $Q_{st}$ estimates for morphological, growth, and metabolic traits in *Salvelinus alpinus*. Between-population variance ($V_b$), family-within-population variance ($V_f$) and within-population variance ($V_w$) are reported. For wild adults, $P_{st}$ are presented across different assumed values of the ratio of between to total additive genetic variance ($\frac{c}{h^2}$). For juveniles from the common garden, $Q_{st}$ was directly estimated. Significant $P_{st}$ and $Q_{st}$ estimates are indicated in bold.

| | $V_b$ | | $V_w$ | | Metric | *p-value* |
|---|---|---|---|---|---|---|
| *Adult* | | | | $\frac{c}{h^2}$ | $P_{st}$ | |
| | | | | 0.33 | **0.45** | **< 0.001** |
| Morphological traits PC1 | 0.005 | - | 0.001 | 0.5 | **0.55** | **< 0.001** |
| | | | | 1 | **0.71** | **< 0.001** |
| | | - | | 0.33 | 0.02 | 0.29 |
| Morphological traits PC2 | $1.91\times10^{-4}$ | - | 0.001 | 0.5 | 0.04 | 0.29 |
| | | - | | 1 | 0.07 | 0.28 |
| | | - | | 0.33 | **0.63** | **< 0.001** |
| Growth PC1 | 5.277 | - | 0.510 | 0.5 | **0.72** | **< 0.001** |
| | | - | | 1 | **0.84** | **< 0.001** |
| | | - | | 0.33 | 0.00 | 0.39 |
| Growth PC2 | $9.75\times10^{-16}$ | - | 0.755 | 0.5 | 0.00 | 0.32 |
| | | - | | 1 | 0.00 | 0.24 |
| *Juvenile* | | $V_f$ | | | $Q_{st}$ | |
| Morphological traits PC1 | $5.9\times10^{-25}$ | $8.1\times10^{-20}$ | 0.001 | | 0.00 | **-** |
| Morphological traits PC2 | $1.72\times10^{-5}$ | $1.63\times10^{-5}$ | $4.23\times10^{-4}$ | | **0.21** | **0.01** |
| Specific growth rate | < 0.001 | 0.002 | 0.004 | | 0.04 | 0.26 |
| Routine metabolic rate | 0.005 | 0.006 | 0.111 | | **0.19** | **0.01** |
| Routine metabolic rate × body mass | 0.151 | 0.005 | 0.010 | | **0.88** | **< 0.001** |
| Routine metabolic rate × temperature | 0.000 | < 0.001 | < 0.001 | | 0.00 | - |

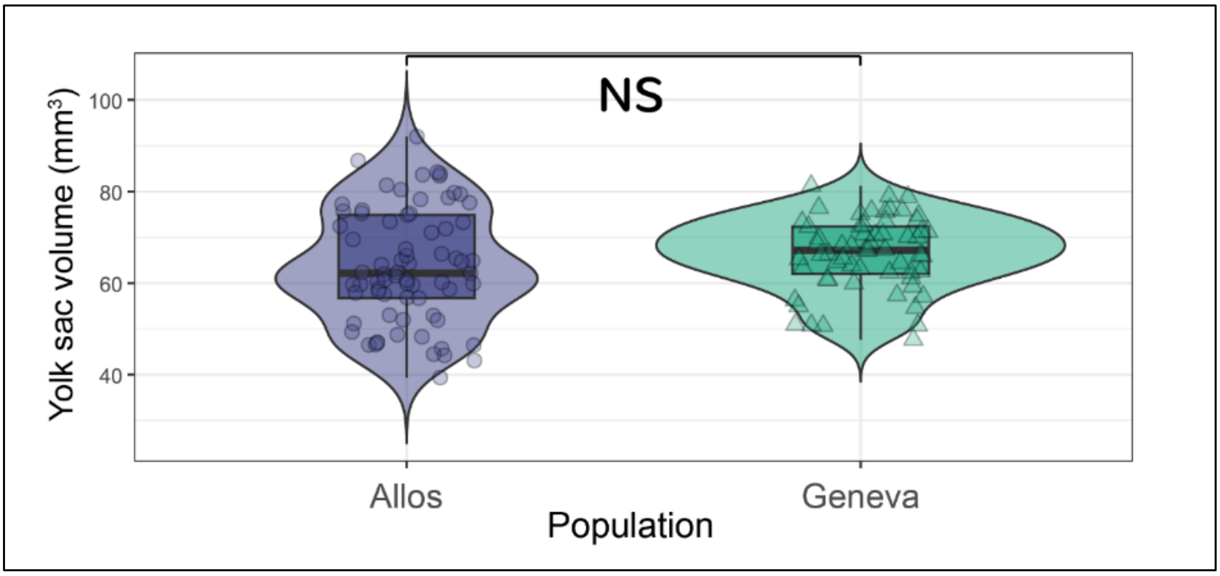


Figure S5. Distribution of yolk sac volume (mm$^3$) in Arctic charr (*Salvelinus alpinus*) from Lake Allos (blue, n = 73; 8 full-sib families) and Lake Geneva (green, n = 60; 6 full-sib families) populations reared under common garden conditions. Yolk sac volume was measured at the hatching stage. No significant difference (NS) in yolk sac volume was detected between populations ($\chi^2 = 0.20$, df = 1, p = 0.655), based on an ANOVA applied to a linear mixed-effects model including family as a random effect.

Table S7. Neutral genetic diversity at fifteen microsatellite loci and pairwise differentiation in Arctic charr (*Salvelinus alpinus*) from Allos and Geneva populations. Number of individuals genotyped (N), observed heterozygosity ($H_o$), expected heterozygosity ($H_e$), inbreeding coefficient ($F_{is}$) and allelic richness ($A_r \pm SD$) are reported for each population. Pairwise $F_{st}$ represents neutral differentiation between populations.

| ***Population*** | ***N*** | $H_o$ | $H_e$ | $F_{is}$ | $A_r \pm SD$ | $F_{st}$ |
|---|---|---|---|---|---|---|
| Allos | 28 | 0.48 | 0.44 | -0.09 | 2.81 ± 0.98 | 0.017 |
| Geneva | 22 | 0.52 | 0.48 | -0.08 | 2.98 ± 1.33 | |

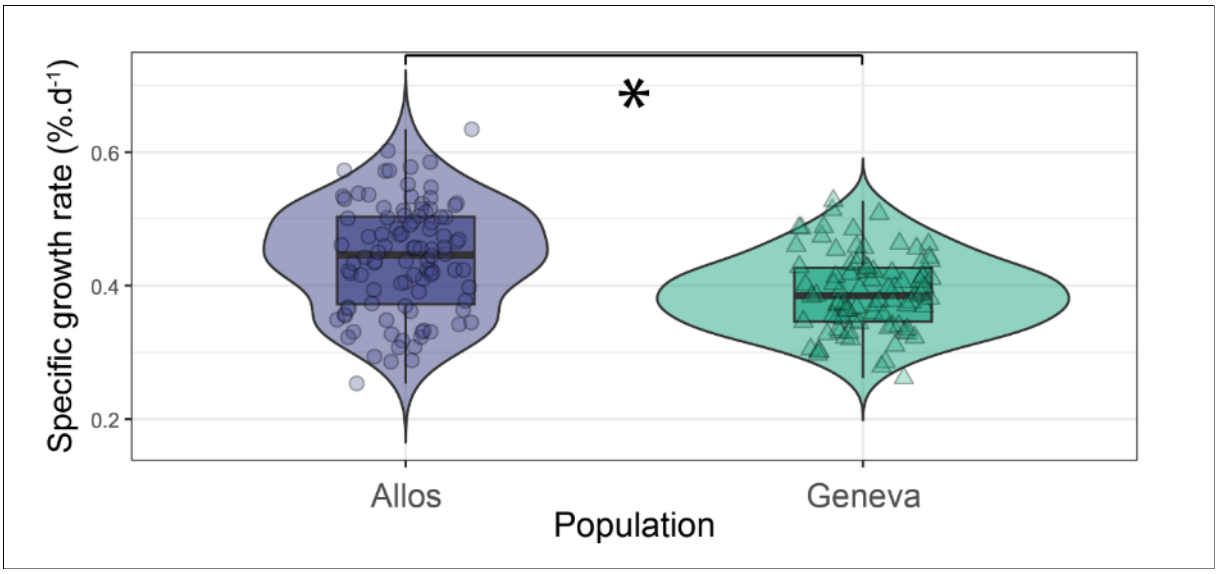


Figure S6. Distribution of Specific Growth Rate (SGR, %.d$^{-1}$) in Arctic charr (*Salvelinus alpinus*) from Lake Allos (blue, n = 100; 10 full-sib families including 2 half-sib families) and Lake Geneva (green, n = 90; 9 full-sib families including 3 half-sib families) populations reared under common garden conditions. Growth rates were measured from hatching to the juvenile stage. Violin plots depict the density distribution of individual SGR values, with embedded boxplots showing the median, interquartile range (IQR), and 1.5×IQR whiskers. Each symbol represents a single individual (Allos, circles; Geneva, triangles). A significant difference in SGR was detected between populations ($\chi^2 = 6.59$, df = 1, p = 0.01), based on an ANOVA applied to a linear mixed-effects model including family as a random effect. Asterisk (*) indicates a significant difference at $\alpha = 0.05$.

### Appendix S1

**A century of allopatry: plasticity and rapid selection shape phenotypic trait variability under contrasting environments.**

Hervé Rogissart [ab*], Martin Daufresne [bc], Guillaume Evanno [d], Jean Guillard [ab], François-Raphael Lubin [ab], Emilie Chancerel [e], Allan Raffard [ab]

[a] Univ. Savoie Mont Blanc, INRAE, CARRTEL, 74200 Thonon-les-Bains, France

[b] Pôle ECLA (OFB, INRAE, USMB), 74200 Thonon-les-Bains, France

[c] INRAE, Aix-Marseille Univ., RECOVER, Aix- en- Provence, France

[d] DECOD (Ecosystem Dynamics and Sustainability), INRAE, Institut Agro, IFREMER, Rennes, France

[e] Université de Bordeaux, INRAE, BIOGECO, Cestas, France

### Appendix S1: Methods of DNA extraction, microsatellite development, genotyping protocol DNA and bioinformatic analysis.

*DNA extraction*

DNA was extracted and isolated using two commercial kits, including the Wizard® SV 96 genomic DNA purification system (Promega) and the QIAwave DNA blood & tissue kit (Qiagen) [1,2]. DNA concentration and quality was measured with a NanoDrop One Spectrophotometer (ThermoFisherScientific).

*Microsatellite development*

DNA from 10 samples were quantified with a Qubit Fluorometer with the Qubit™ dsDNA BR assay kit and pooled in equimolar concentration. The DNA pool was then purified using 1.8 X Agencourt AMPure XP beads (Beckman Coulter, UK). Whole genome DNA library were prepared using Qiagen QIASeq FX DNA library preparation kit and sequenced on an Illumina iseq100 in a 300 bp single-read configuration.

QDD software was used to discover and select microsatellites from the sequencing data [3]. Primer design parameters were set to target 100 pb to 180 pb amplicons and with primer parameters optimized for multiplex PCR [4]. A total of 40 primer pairs (3 tetranucleotide, 9 trinucleotide and 28 dinucleotide motifs) were selected based on criteria maximizing amplification success [5] and polymorphisms (i.e. high number of repeats). Twenty other SSR marker sequences from *Salvelinus alpinus* and in phylogenetically closest species (*Salvelinus fontinalis* and *Salvelinus malma*) were extracted from NCBI database. QDD pipe3 was used to design primers under the same conditions as above (3 tetranucleotide, 7 trinucleotide and 10 dinucleotide motifs).

Standards desalt oligonucleotides were ordered with universal sequence (US) attached to the 5' end of each primer (US for forward primer: ACACTCTTTCCCTACACGACGCTCTTCCGATCT and US for reverse primer: GACTGGAGTTCAGACGTGTGCTCTTCCGATCT). Each primer pair was tested for amplification on a pool of 95 DNA samples. The PCR was performed in a final volume of 10µL using Hot Firepol Blend master mix (Solis Biodyne), 10 ng of DNA and 0.2 µM of each primer. The PCR conditions consisted of an initial denaturation at 95°C for 15-min followed by 35 cycles of denaturation at 95°C for 20 s, annealing at 59°C for 60 s, extension at 72°C for 30 s, and a final extension step at 72°C for 10-min. Amplification was check on a 3% agarose gel. Primer pairs that showed consistent amplification with amplicon at the expect size were integrated in the primer pool for the multiplex PCR (only four primer pairs were excluded at this stage).

*Genotyping protocol*

Multiplex PCR amplification of the 56 selected markers was performed in a final volume of 5 µL using 5X Hot Firepol Multiplex master mix (Solis Biodyne), 0.05 µM of each primer, and 10 ng of DNA. The PCR conditions consisted of an initial denaturation at 95°C for 12-min followed by 35 cycles of denaturation at 95°C for 30 s, annealing at 59°C for 180 s, extension at 72°C for 30 s, and a final extension step at 72°C for 10-min. The sequencing libraries of each mutiplex was constructed using a second PCR that attached adapters and sample-specific pairs of indexes (10 bp unique sequences) to each side of the amplicons by targeting the universal sequence attached to the locus specific primers. The indexing PCR is setup in a volume of 5 µL using 5X Hot Firepol Multiplex master mix (Solis Biodyne), 1,25 µL of amplicon and 0.5 µM of each of the forward and reverse adapters. The PCR conditions consisted in an initial denaturation at 95°C for 12-min followed by 15 cycles of denaturation at 95°C for 30 s, annealing at 59°C for 90 s, extension at 72°C for 30 s, and a final extension step at 72°C for 10-min. Libraries are then pooled, purified with 1.2x SPRI magnetic beads and quantified with Qubit™ dsDNA BR assay kit. A reconditioning PCR [6] was performed on 400 ng of the pool with 0.5 µM Illumina P5 (5'-AATGATACGGCGACCACCGAGATCT-3') and P7 (5'-CAAGCAGAAGACGGCATACGAGAT-3') PCR primers and 10 µL of 5X Hot Firepol Multiplex master mix. The condition for PCR amplification, was as follows: initial denaturation at 95°C for 12-min, followed by 1 amplification cycle of 30 sec at 95°C, 90 sec at 59°C and 30 sec at 72°C, followed by final extension at 720°C for 10-min. An ultime purification with 0.95x SPRI magnetic beads was performed. We checked the library pool quality on a Tapestation 4200 (Agilent Technologies, Santa Clara, CA) and quantified it using NEBNext Library Quant Kit (New England Biolabs) on a Mic qPCR Cycler (Bio Molecular Systems).

The whole procedure was performed on 95 samples and a negative control (containing water instead of DNA), and independently repeated once on all samples in order to 1) optimize the bioinformatic pipeline to each locus, and 2) estimate locus-level allelic error rate (number of allele mismatches between replicates divided by the total number of alleles compared), and finally 3) select loci that produced repeatable genotypes for the final genotypic dataset. The sequencing was done on an Illumina iseq100 or NextSeq 2000 2x150 pb paired-end configuration.

*Bioinformatic analysis*

The analysis protocol consists of three main steps explained in [4]: (1) sequence preparation, (2) preliminary marker validation analysis by comparing replicates and testing different analysis parameters and (3) final genotyping analysis using the optimal parameters on the validated markers.

Table S1: Characteristics of the microsatellite loci used for genetic analyses in *Salvelinus alpinus*. For each locus, the table reports the locus name, microsatellite motif, amplicon size, forward (F) and reverse (R) primer sequences; mean and standard deviation of per-locus per-sample coverage; analysis strategy and parameter set used in FDSTools; number of alleles detected based on sequence data (NallelesSequence) and on allele size (NalleleSize); as well as the proportion of missing data (MissingRate) and estimated allelic error (AllelicError). Loci that failed to amplify during simplex PCR (n.s.#) were not sequenced.

| Locus | Read ID (+de novo; *from NCBI) | Repeat motif | Amplicon size | Forward [a] and Reverse [b] primers sequence (5'-3') | Mean coverage locus/sample [c] | SD coverage | Analysis strategy [d] | Parameter set [e] | N alleles Sequence | N alleles Size | Missing Rate | Allelic Error |
|---|---|---|---|---|---|---|---|---|---|---|---|---|
| SSRSEQ_iServ_UDI_001 | +FS10001056:97:BTR99620-0718:1:1101:10670:2450 | (CAGA)6 | 129 | F: TCTTTGTCTCCGGGTGATATTCTGCT<br>R: TGGAATTCAGACGTGTGCTCTTCCGA | n.s. | n.s. | F | | | | | |
| SSRSEQ_iServ_UDI_002 | +FS10001056:97:BTR99620-0718:1:1101:11690:2830 | (GTTA)6 | 155 | F: CAGGCCTGCGTACGGTGATGATGT<br>R: GCAGGGAGGAGAGGACAGACTAGCA | 1979,99 | 734,43 | FL | PS1 | 2,00 | 1,00 | 0,01 | 0,01 |
| SSRSEQ_iServ_UDI_003 | +FS10001056:97:BTR99620-0718:1:1101:12900:2130 | (TGTC)10 | 180 | F: AGGGCACAGAGGAAACAGGCTACCT<br>R: GCTTTGAGGAAGGGCAGAAGGCTG | 4375,81 | 1544,76 | FL | PS1 | 6,00 | 6,00 | 0,01 | 0,00 |
| SSRSEQ_iServ_UDI_004 | +FS10001056:97:BTR99620-0718:1:1101:13800:1740 | (TGT)6 | 145 | F: GCAGATATTAGTTCGGGCAGTGAGGC<br>R: CACAGCAGCCTAGAAGCTATGCACA | 4,52 | 3,20 | F | | | | | |
| SSRSEQ_iServ_UDI_005 | +FS10001056:97:BTR99620-0718:1:1101:2450:1370 | (AAG)11 | 119 | F: TTCTGTACTATGGCCGAGACATCCC<br>R: TGGCATCCAACCTTCCTCGTTGACC | 7046,78 | 2419,90 | FL | PS1 | 4,00 | 4,00 | 0,01 | 0,00 |
| SSRSEQ_iServ_UDI_006 | +FS10001056:97:BTR99620-0718:1:1101:3670:1370 | (CTA)6 | 180 | F: AGGCCTGTGTTTGACGTTGCAAGTCA<br>R: TGACCGTAGTGAAGACCAGCGCAAA | 133,85 | 70,23 | F | | | | | |
| SSRSEQ_iServ_UDI_007 | +FS10001056:97:BTR99620-0718:1:1103:13350:1810 | (TTG)7 | 100 | F: CATGGTTGTTCCTGTTGGGTAGCAA<br>R: TGCTCTTCCGATCTGAAGTTGCTAC | 285,69 | 115,67 | F | | | | | |
| SSRSEQ_iServ_UDI_008 | +FS10001056:97:BTR99620-0718:1:1104:8350:3440 | (CCT)7 | 100 | F: AAACCCTCTGAGATATTCCTCCCTCC<br>R: ACGTCTTCCTCCCTTACACCTCGC | 4831,25 | 1805,45 | FL | PS1 | 1,00 | 1,00 | 0,01 | 0,00 |
| SSRSEQ_iServ_UDI_009 | +FS10001056:97:BTR99620-0718:1:1110:8170:2450 | (TCA)7 | 100 | F: CCAGCAGAGTGATAGCTTTCTTCCG<br>R: AAGGTGGTAGAAGCGTTTGATGAGGG | 8902,16 | 3032,01 | FL | PS1 | 2,00 | 2,00 | 0,01 | 0,00 |
| SSRSEQ_iServ_UDI_010 | +FS10001056:97:BTR99620-0718:1:1113:12820:1200 | (ACA)7 | 100 | F: CAACAAGCTGGTGAAGTTGCTACCC<br>R: GTCATGGTAGTTTCTGTTGGGAAGCA | 1379,09 | 488,12 | F | | | | | |
| | +FS10001056:97:BTR99620-0718:1:1114:9460:3550 | (GAT)8 | 100 | F: GTTGGAGATCCTGGGACCAGTACCA | 8065,90 | 2741,05 | FL | PS1 | 3,00 | 3,00 | 0,01 | 0,01 |

| | | | | | | | | | | | | | |
|---|---|---|---|---|---|---|---|---|---|---|---|---|---|
| SSRSEQ_iServ_UDI_011 | | | | R: | GATCAGTCGGAGTCATCCTTGTCGC | | | | | | | | |
| SSRSEQ_iServ_UDI_012 | +FS10001056:97:BTR99620-0718:1:1116:9930:2610 | (CCA)6 | 100 | F:<br>R: | CGTTGTCCGTAGCTTATACCTGCTC<br>GGTTTGCTGATGTCAACGTTGTGAA | 5212,91 | 1883,10 | F | | | | | |
| SSRSEQ_iServ_UDI_013 | +FS10001056:97:BTR99620-0718:1:1101:1140:1310 | (GT)18 | 164 | F:<br>R: | AGCCGCCGAGTCATGATACACTCAA<br>TTGTGCCCTGTAGGTTGTTGCCACC | 2388,45 | 1247,11 | F | | | | | |
| SSRSEQ_iServ_UDI_014 | +FS10001056:97:BTR99620-0718:1:1101:14240:3000 | (AC)19 | 138 | F:<br>R: | GGGACATGACCTCTCAGTGCCACAGT<br>ACCTCCGAGTCCTTTGCAGTCACAT | 1911,42 | 1211,30 | RF | PS1 | 8,00 | 8,00 | 0,08 | 0,01 |
| SSRSEQ_iServ_UDI_015 | +FS10001056:97:BTR99620-0718:1:1101:15770:2510 | (AC)21 | 121 | F:<br>R: | GGGCTCTGGATGAAGAGAAGAGATT<br>TGGTACGTGTGACACTCCTGATGTG | 40,12 | 19,93 | F | | | | | |
| SSRSEQ_iServ_UDI_016 | +FS10001056:97:BTR99620-0718:1:1101:16480:1360 | (TG)10 | 176 | F:<br>R: | CCACAGACAGAAGAACATGCGGCGT<br>GTGTGCTCTTCCGATCTGCCAAGCG | n.s. | n.s. | F | | | | | |
| SSRSEQ_iServ_UDI_017 | +FS10001056:97:BTR99620-0718:1:1101:1700:3540 | (GT)30 | 132 | F:<br>R: | GTTACTGGCCCAACGCTCTAACCGC<br>TGGGACCCAGCGTTAGAGGTGTGAC | 2,53 | 2,45 | F | | | | | |
| SSRSEQ_iServ_UDI_018 | +FS10001056:97:BTR99620-0718:1:1101:4690:2730 | (AC)13 | 137 | F:<br>R: | CCACTGATCTCTTCTTCCAGAGGCC<br>TGTGCTTCGCATTGTGTTAGTTGC | 5826,90 | 2245,39 | FL | PS1 | 5,00 | 4,00 | 0,01 | 0,01 |
| SSRSEQ_iServ_UDI_019 | +FS10001056:97:BTR99620-0718:1:1101:5480:1400 | (CA)19 | 165 | F:<br>R: | GGCAAGGATGTCACACCGCGGATTC<br>GCGATGGTGCGAAAGATTCTCGTGT | 0,15 | 0,65 | F | | | | | |
| SSRSEQ_iServ_UDI_020 | +FS10001056:97:BTR99620-0718:1:1101:7070:2390 | (GT)17 | 147 | F:<br>R: | TGCTGACCTTTGACCCTTCGATCAGA<br>TCGAAGTGGTGTCCTTCAGGTGTGA | 0 | 0 | F | | | | | |
| SSRSEQ_iServ_UDI_021 | +FS10001056:97:BTR99620-0718:1:1101:9890:3590 | (TG)26 | 113 | F:<br>R: | GCCAACATTCCAGCGGAACCCACAA<br>ACCCTAGTGCCATCAGCTACTGGGT | 275,82 | 149,16 | F | | | | | |
| SSRSEQ_iServ_UDI_022 | +FS10001056:97:BTR99620-0718:1:1102:12090:1010 | (GT)10 | 153 | F:<br>R: | AAAGTAGGCCTTCCTCAGCGTGGCT<br>GCTCCTGCCAGCTGGGTCCTTATTG | 8551,21 | 2954,59 | FL | PS1 | 2,00 | 2,00 | 0,01 | 0,01 |
| SSRSEQ_iServ_UDI_023 | +FS10001056:97:BTR99620-0718:1:1102:13060:1330 | (GT)11 | 136 | F:<br>R: | CCGTGCATCGACAGGACAGAACAGC<br>ACCTCAGGCGAATAGTACCTCGAGAA | 4206,21 | 1557,38 | F | | | | | |
| SSRSEQ_iServ_UDI_024 | +FS10001056:97:BTR99620-0718:1:1102:14600:1600 | (GT)13 | 180 | F:<br>R: | TGTCACTGCCACCAAGACCACTGGA<br>TCCGATCTCACATGGTTTGGTGGCT | 4088,65 | 2012,47 | RF | PS1 | 5,00 | 5,00 | 0,02 | 0,00 |

| | | | | | | | | | | | | |
|---|---|---|---|---|---|---|---|---|---|---|---|---|
| SSRSEQ_iServ_UDI_025 | +FS10001056:97:BTR99620-0718:1:1102:3300:1160 | (GT)25 | 122 | F: AGTGAAGCTGTTGTAGGTGGGCAGT<br>R: GCCCAGCTCCTCTTGTTCTTTCTCGC | 4699,26 | 1726,03 | RF | PS2 | 5,00 | 5,00 | 0,01 | 0,01 |
| SSRSEQ_iServ_UDI_026 | +FS10001056:97:BTR99620-0718:1:1102:4800:1390 | (AC)15 | 129 | F: CCCTGCTGGCTTCCTTTCCTTCTCA<br>R: TACACGCCTAGACCCTCAATGTGGT | 7083,16 | 2544,29 | F | | | | | |
| SSRSEQ_iServ_UDI_027 | +FS10001056:97:BTR99620-0718:1:1102:6940:1630 | (CA)25 | 140 | F: ATGGAAGCTGGGTCAAAGAGGCGGA<br>R: CTCTGTGTCTCACTGCCCGTGT | 2,03 | 2,04 | F | | | | | |
| SSRSEQ_iServ_UDI_028 | +FS10001056:97:BTR99620-0718:1:1102:8760:1640 | (TG)14 | 142 | F: GTGTCCCTACAACCTGAGGAAAGAA<br>R: AGCATTTCACTGTTGGTCTCTGCCT | 5708,45 | 2657,35 | FL | PS2 | 2,00 | 2,00 | 0,01 | 0,00 |
| SSRSEQ_iServ_UDI_029 | +FS10001056:97:BTR99620-0718:1:1102:9760:1680 | (TG)23 | 116 | F: AGGAGGAAACCCACTGCTGAGCGAA<br>R: TGGGTATGGTGAGAACAGCTTGTTGG | n.s. | n.s. | F | | | | | |
| SSRSEQ_iServ_UDI_030 | +FS10001056:97:BTR99620-0718:1:1104:12240:1720 | (GT)15 | 100 | F: CTACTGCCACCGCATGACATTGTAG<br>R: GTGACTGGAGTTCAGACGTGTGCTC | n.s. | n.s. | F | | | | | |
| SSRSEQ_iServ_UDI_031 | +FS10001056:97:BTR99620-0718:1:1104:14650:1970 | (AC)9 | 100 | F: AGCGGAACTGGCATTTGGGTATGT<br>R: AGGGAGAGTGGCATGGCATTTAGGA | 6297,16 | 2196,35 | FL | PS2 | 3,00 | 2,00 | 0,01 | 0,00 |
| SSRSEQ_iServ_UDI_032 | +FS10001056:97:BTR99620-0718:1:1105:11220:1100 | (TG)10 | 100 | F: CTTTGTCCTTGGAACTCCCGCCTG<br>R: CAACCCAGACCGGCATGGCACTCAA | 5066,41 | 1824,51 | F | | | | | |
| SSRSEQ_iServ_UDI_033 | +FS10001056:97:BTR99620-0718:1:1108:6330:2350 | (AC)9 | 100 | F: TCTGGTGACTTGTCAGGACGTTCTG<br>R: GCAAAGGTTCCATGTCCACTGTCCC | 5780,85 | 2052,15 | F | | | | | |
| SSRSEQ_iServ_UDI_034 | +FS10001056:97:BTR99620-0718:1:1108:15650:2300 | (CA)12 | 100 | F: GAAGTCCTACGACAGGCTGCTACAGG<br>R: GTGCCGTTTGATGCCATGTGGT | 4764,15 | 1660,16 | FL | PS2 | 4,00 | 4,00 | 0,04 | 0,00 |
| SSRSEQ_iServ_UDI_035 | +FS10001056:97:BTR99620-0718:1:1109:13720:1600 | (AG)10 | 100 | F: GCCATACTGGAGCACATGGGCAGATT<br>R: TCAGACAGTCAACACCTCCATAGCT | 5459,83 | 1869,32 | FL | PS1 | 3,00 | 3,00 | 0,01 | 0,01 |
| SSRSEQ_iServ_UDI_036 | +FS10001056:97:BTR99620-0718:1:1109:15630:2670 | (GT)12 | 100 | F: CGTCCACAAACTGGCCACTGCTCTA<br>R: GCTGTGATGGTCTGATGCCAAGCCC | 4877,50 | 1715,84 | FL | PS1 | 3,00 | 2,00 | 0,01 | 0,00 |
| SSRSEQ_iServ_UDI_037 | +FS10001056:97:BTR99620-0718:1:1109:5410:2210 | (TG)9 | 100 | F: CCGAGTGGGAGGACATGGAGTTTGT<br>R: ACTCAGTGCCAAAGACATAAGAGCC | 6574,70 | 2320,11 | FL | PS1 | 3,00 | 3,00 | 0,16 | 0,01 |

| | | | | | | | | | | | | | |
|---|---|---|---|---|---|---|---|---|---|---|---|---|---|
| SSRSEQ_iServ_UDI_038 | +FS10001056:97:BTR99620-0718:1:1110:2450:1440 | (TG)14 | 100 | F:<br>R: | TACTGCTTCTGTCCTTAAACCAGCCG<br>TCAGGCACCGCAGAGACAGACACA | 839,87 | 712,48 | FL | PS1 | 7,00 | 7,00 | 0,31 | 0,01 |
| SSRSEQ_iServ_UDI_039 | +FS10001056:97:BTR99620-0718:1:1112:9020:1840 | (TC)15 | 100 | F:<br>R: | AATGGCTAGACATCTCCCTCAAGCT<br>TGTCAAGCCTCTTGATTGCACGGAT | 5801,14 | 2009,87 | FL | PS1 | 6,00 | 6,00 | 0,01 | 0,00 |
| SSRSEQ_iServ_UDI_040 | +FS10001056:97:BTR99620-0718:1:1112:6080:1930 | (GA)14 | 100 | F:<br>R: | CACAAAGCCCGGCAAGATAGG<br>CTGGCAGTGCTCGTAACAAAC | 4494,35 | 1654,91 | FL | PS1 | 6,00 | 5,00 | 0,01 | 0,01 |
| SSRSEQ_iServ_UDI_041 | *AF129788.1 | (AC)10 | 156 | F:<br>R: | GAACATGGCCCTGGACAGCAGGAAC<br>TCACCAGGTCCCGCATCGCATACTT | 1807,45 | 660,74 | FL | PS1 | 3,00 | 3,00 | 0,01 | 0,01 |
| SSRSEQ_iServ_UDI_042 | *AF537304.1 | (CA)33 | 123 | F:<br>R: | CCCTCCCTTCCGCTTTCACACAGAC<br>TTACGTGAAAGCTACCTTGCGTGCA | 330,03 | 163,41 | F | | | | | |
| SSRSEQ_iServ_UDI_043 | *AF537306.1 | (AC)12 | 142 | F:<br>R: | CCTCCCTGACTCCATGGACTACGACA<br>TGTGTGCGTTCGCGTGTTCTTATGC | 4445,61 | 2054,43 | FL | PS1 | 4,00 | 4,00 | 0,01 | 0,00 |
| SSRSEQ_iServ_UDI_044 | *AY168186.1 | (GTGT)10 | 149 | F:<br>R: | TGGTCTGTCTATCTAAAGAGGCTGT<br>TGTCTTGGTGATTTCAGAGCTCTGT | 1468,51 | 728,37 | RF | PS1 | 3,00 | 3,00 | 0,01 | 0,03 |
| SSRSEQ_iServ_UDI_045 | *AY168188.1 | (TGTG)10 | 173 | F:<br>R: | AAGGGTTGAATAGAAGAGCTGCCTA<br>TCCCTCAGCTTCAAAGACACTCACA | 17,30 | 9,73 | F | | | | | |
| SSRSEQ_iServ_UDI_046 | *AY168197.1 | (ATAG)17 | 174 | F:<br>R: | AGGTAGAGCATGAGCAGCGAGAAGA<br>TTGTTGTCTTCCCTCGGTCTCTCCC | 327,07 | 184,20 | FL | PS1 | | | | |
| SSRSEQ_iServ_UDI_047 | *MN530969.1 | (TC)22 | 173 | F:<br>R: | CCCATGGCTGTTCACCACATAGAGT<br>TCCTCCCATCACACATGGTATTGCT | 11396,46 | 3970,32 | FL | PS1 | 3,00 | 2,00 | 0,01 | 0,01 |
| SSRSEQ_iServ_UDI_048 | *AF537307.1 | (TG)25 | 143 | F:<br>R: | CCTGAGATGAAACCAGGGCCGCAAC<br>AGACACCCTAACACCCTTGGTTCTGT | 1565,45 | 893,73 | F | | | | | |
| SSRSEQ_iServ_UDI_049 | *AF537312.1 | (CA)18 | 124 | F:<br>R: | TTCATATGCTCTGCCGCGCACGTTT<br>TATTAACGCCCACTCCCTGGAGACA | 153,79 | 107,83 | F | | | | | |
| SSRSEQ_iServ_UDI_050 | *AY168189.1 | (ATG)6 | 129 | F:<br>R: | TGGGCCACTGTTGTGTTGCTTTGGT<br>TAACTAGTAGCTGTAACCCTGCTGC | 6168,38 | 2191,56 | FL | PS1 | 5,00 | 3,00 | 0,01 | 0,01 |
| | *AY168190.1 | (TGA)7 | 159 | F: | GGTCTCGGTCAACATCAGCCTCACT | 2,60 | 2,55 | F | | | | | |

| | | | | | | | | | | | | |
|---|---|---|---|---|---|---|---|---|---|---|---|---|
| SSRSEQ_iServ_UDI_051 | | | | R: GGTGTCAATTTAGGAACTGCTAGCCC | | | | | | | | |
| SSRSEQ_iServ_UDI_052 | *AY168191.1 | (TCA)8 | 171 | F: CTTCCCTGTTCGCTGCAGACCGATG<br>R: CCAGAGTTGTTCCTGGCCTGGTCCA | 4943,38 | 1819,85 | FL | PS1 | 1,00 | 1,00 | 0,01 | 0,00 |
| SSRSEQ_iServ_UDI_053 | *AY168192.1 | (GAT)16 | 156 | F: TCTGGGTAATGTAGTCTCTGGTGGG<br>R: ATGTCTGAACCCTGATTGTGAACGT | 5820,13 | 2298,69 | RF | PS1 | 2,00 | 2,00 | 0,01 | 0,00 |
| SSRSEQ_iServ_UDI_054 | *AY168193.1 | (TGA)13 | 126 | F: GGGAGCCCAGACTATATTGACGATGA<br>R: CCAGATGCTGTCCTTGTTGCCACCG | 9493,60 | 3231,86 | FL | PS1 | 4,00 | 4,00 | 0,01 | 0,01 |
| SSRSEQ_iServ_UDI_055 | *AY168195.1 | (TGA)9 | 148 | F: TTGGGTCGTGGTGCAGGCACTAACT<br>R: GCAACACGTCCTCCTCTTCCAGGGA | 9269,03 | 3275,77 | FL | PS1 | 2,00 | 1,00 | 0,01 | 0,01 |
| SSRSEQ_iServ_UDI_056 | *AY327124.1 | (AC)11 | 154 | F: TTCACATTTCCTGTTGCTGAAGGAT<br>R: ACTTGTTCTCCTGCCAGCCAACAGT | 1332,48 | 497,08 | FL | PS1 | 2,00 | 1,00 | 0,01 | 0,00 |
| SSRSEQ_iServ_UDI_057 | *AY327127.1 | (CA)29 | 152 | F: ACGCTGCTGTTGAGAATGCCAACCT<br>R: GGTCTACTGTTAGATTAGATGCGCG | 2914,58 | 1148,01 | RF | PS1 | 3,00 | 3,00 | 0,15 | 0,00 |
| SSRSEQ_iServ_UDI_058 | *AY168187.1 | (TGA)5 | 105 | F: CCACATAATAGAACACTGGAGCCGT<br>R: TGTGTCTAAAGATCACAGAGATGGG | 2485,77 | 909,72 | RF | PS1 | 3,00 | 3,00 | 0,01 | 0,01 |
| SSRSEQ_iServ_UDI_059 | *AY168194.1 | (CA)7 | 117 | F: GAACAGCCATCACACCATCTCCTGT<br>R: ACGTGTCTGAGTGTATATGTGAGTAC | 3079,62 | 1111,76 | FL | PS1 | 1,00 | 1,00 | 0,01 | 0,00 |
| SSRSEQ_iServ_UDI_060 | *AY327125.1 | (CA)7 | 160 | F: TGGTTAGCCACATCACACTGATCAG<br>R: TCTTGACCTCACTCTGTTCAGAGCA | 273,85 | 172,37 | F | | | | | |
| **Mean** | | | | | **3647** | **1368** | | | **4** | **3** | **0,029** | **0,004** |
| **Overall** | | | | | | | | | **116** | **106** | | |

[a], the universal Illumina adapter overhang sequences ACACTCTTTCCCTACACGACGCTCTTCCGATCT was added to the 5’ end of the forward primers;
[b], the universal Illumina adapter overhang sequences GACTGGAGTTCAGACGTGTGCTCTTCCGATCT was added to the 5’ end of the reverse primers;
[c] loci that did not amplify during the test simplex PCR were not sequenced (n.s.), 0 indicate locus that amplified in simplex PCR but not in the multiplex PCR;
[d], F: Failed, FL: Full Length, RF: Repeat Focused;
[e], FDSTools parameters as followed: PS1 corresponds to -s-1=50,+1=10 -m=15 -n=20 and PS2 corresponds to -s-1=70,+1=10 -m=10 -n=20.
